# Convergent evolution of viral-like Borg archaeal extrachromosomal elements and giant eukaryotic viruses

**DOI:** 10.1101/2024.11.05.622173

**Authors:** Jillian F. Banfield, Luis E. Valentin-Alvarado, Ling-Dong Shi, Colin Michael Robinson, Rebecca S. Bamert, Fasseli Coulibaly, Zachary K. Barth, Frank O. Aylward, Marie C Schoelmerich, Shufei Lei, Rohan Sachdeva, Gavin J. Knott

**Affiliations:** Biomedicine Discovery Institute, Monash University; Innovative Genomics Institute, UC Berkeley; Earth and Planetary Science, UC Berkeley; Environmental Science, Policy and Management, UC Berkeley; Plant and Microbial Biology, UC Berkeley; Virginia Polytechnic Institute and State University.

## Abstract

Borgs are huge extrachromosomal elements of anaerobic methane-oxidizing archaea. They exist in exceedingly complex microbiomes, lack cultivated hosts and have few protein functional annotations, precluding their classification as plasmids, viruses or other. Here, we used *in silico* structure prediction methods to investigate potential roles for ∼10,000 Borg proteins. Prioritizing analysis of multicopy genes that could signal importance for Borg lifestyles, we uncovered highly represented de-ubiquitination-like Zn-metalloproteases that may counter host targeting of Borg proteins for proteolysis. Also prevalent are clusters of multicopy genes for production of diverse glycoconjugates that could contribute to decoration of the host cell surface, or of putative capsid proteins that we predict multimerize into hexagonal arrays. Features including megabase-scale linear genomes with inverted terminal repeats, genomic repertoires for energy metabolism, central carbon compound transformations and translation, and pervasive direct repeat regions are shared with giant viruses of eukaryotes, although analyses suggest that these parallels arose via convergent evolution. If Borgs are giant archaeal viruses they would fill the gap in the tri(um)virate of giant viruses of all three domains of life.

**One Sentence Summary:** Protein analyses, informed by *in silico* protein structure prediction, revealed that Borgs share numerous features with giant eukaryotic viruses, suggesting that Borgs have a viral-like lifestyle and evolutionary convergence of large extrachromosomal elements across the Domains of Life.

## MAIN TEXT

Assigning function to proteins of extrachromosomal elements (ECEs) of Archaea, the third Domain of life, is challenging (Prangishvili et al. 2017). This is likely due to historical bias towards the characterization of proteins from model organisms that do not represent the staggering sequence diversity acquired over billions of years of evolution. Lack of understanding about archaeal ECEs is important, given that Archaea play key roles in the methane, nitrogen and carbon cycles. Further, as Archaea are the likely ancestors to Eukaryotes (Eme et al. 2023), their ECEs may illuminate the evolutionary origin of eukaryotic viruses. Of particular interest to us are Borgs, huge ECEs that replicate in anaerobic methane-oxidizing archaea (Al-Shayeb et al. 2022). Borgs and their host *Methanoperedens* archaea each account for only a miniscule fraction of the DNA in the soil in which they occur. *Methanoperedens* have not yet been obtained in pure culture and all laboratory *Methanoperedens* enrichments lack Borgs. Thus, all information about Borgs must be acquired from their nucleic acid sequences. Their linear genomes range up to 1.1 Mbp in length and are terminated by long inverted repeats. These features are shared by some plasmids (Wagenknecht et al. 2010), and it has been stated that Borgs are (obviously) plasmids. However, linear genomes are also typical of some viruses of archaea (Krupovic et al. 2018) and many giant eukaryotic viruses (Talbert et al. 2023). Our recent study used *in silico* structure prediction to uncover the presence of some capsid-like proteins (Al-Shayeb et al. 2022; Schoelmerich et al. 2024), raising the possibility that Borgs may be virus-like.

Standard annotation methods predict functions for ∼20% of Borg proteins, yet reveal a surprising inventory of metabolic and energy-relevant capacities, encoded by both single and multi-gene clusters (Al-Shayeb et al. 2022). Examples include four proteins of the methyl-coenzyme M reductase (MCR) complex central to methane metabolism, two proteins for biosynthesis of F430 (the cofactor for MCR), electron transfer via ubiquitous multiheme cytochromes, polyhydroxybutyrate production, and nitrogen fixation (Al-Shayeb et al. 2022; Schoelmerich et al. 2024). Reasoning that existence of multi-copy proteins in Borg genomes could indicate their particular importance, we investigated proteins from the subfamilies most highly represented in Borg genomes using *in silico* structure prediction. As analyses that rely on gene content require accurate and complete or very nearly complete genomes, we focused on the predicted proteomes of the 12 complete and 5 essentially complete Borg genomes that we published recently (Al-Shayeb et al. 2022; Schoelmerich et al. 2024). We place the findings into context via an analysis of proteome content and organization of all 17 Borg genomes.

### Borg genomes encode multicopy proteins, many of which are related to deubiquitinases

Structures of >10,000 proteins were predicted using AlphaFold2 (Jumper et al. 2021; Mirdita et al. 2022) and AlphaFold3 (Abramson et al. 2024). This set included all proteins of seven Borgs (Orange, Black, Green, Amber, Amethyst, Cobalt and Ruby; **Supplementary Data 1A**) from the previously defined two major clades (Schoelmerich et al. 2024) and a subset of proteins from other Borgs (**Supplementary Data item 1B**). Output models were assessed in terms of prediction confidence (**Supplementary Data item 1C**) and possible functions suggested based on homology to existing experimental structures in the Protein Data Bank (PDB; **Table S1-S7**) using FoldSeek (van Kempen et al. 2023) or Dali (Holm 2022). Confident predictions (plDDT > 0.7) were generated for 63% of proteins, of which 36% had potentially informative hits in PDB (bitscores > 200; **Figure S1**).

We identified 28 Borg protein subfamilies (clusters of sequences with similarities high enough to suggest shared functions) that have an average of two or more representatives in 17 genomes (**Figure 1A**, **Table S8)**. The individual subfamilies were defined previously (Schoelmerich et al. 2024). Most of the 1596 proteins in the 28 subfamilies had no, or only poorly defined, functions assigned using standard annotation methods. The most highly represented (subfam1001) had a few low scoring structural matches to proteins in the PDB. The third most highly represented, subfam0238, had similar functional predictions, as did representatives of three much less commonly detected subfamilies (0196, 0759, 1928) and a few structurally similar proteins not assigned to any subfamily. The 315 proteins are phylogenetically intermixed (**Figure 1B, Supplementary Data 2**), thus all were assigned to Group 1, with an average of 18.5 representatives per genome. The structure predicted using an alignment of the 315 protein sequences without using PDB templates was closely similar to that predicted using PDB templates and had similar fold confidence scores (**Figure S2**).

**Figure 1:**
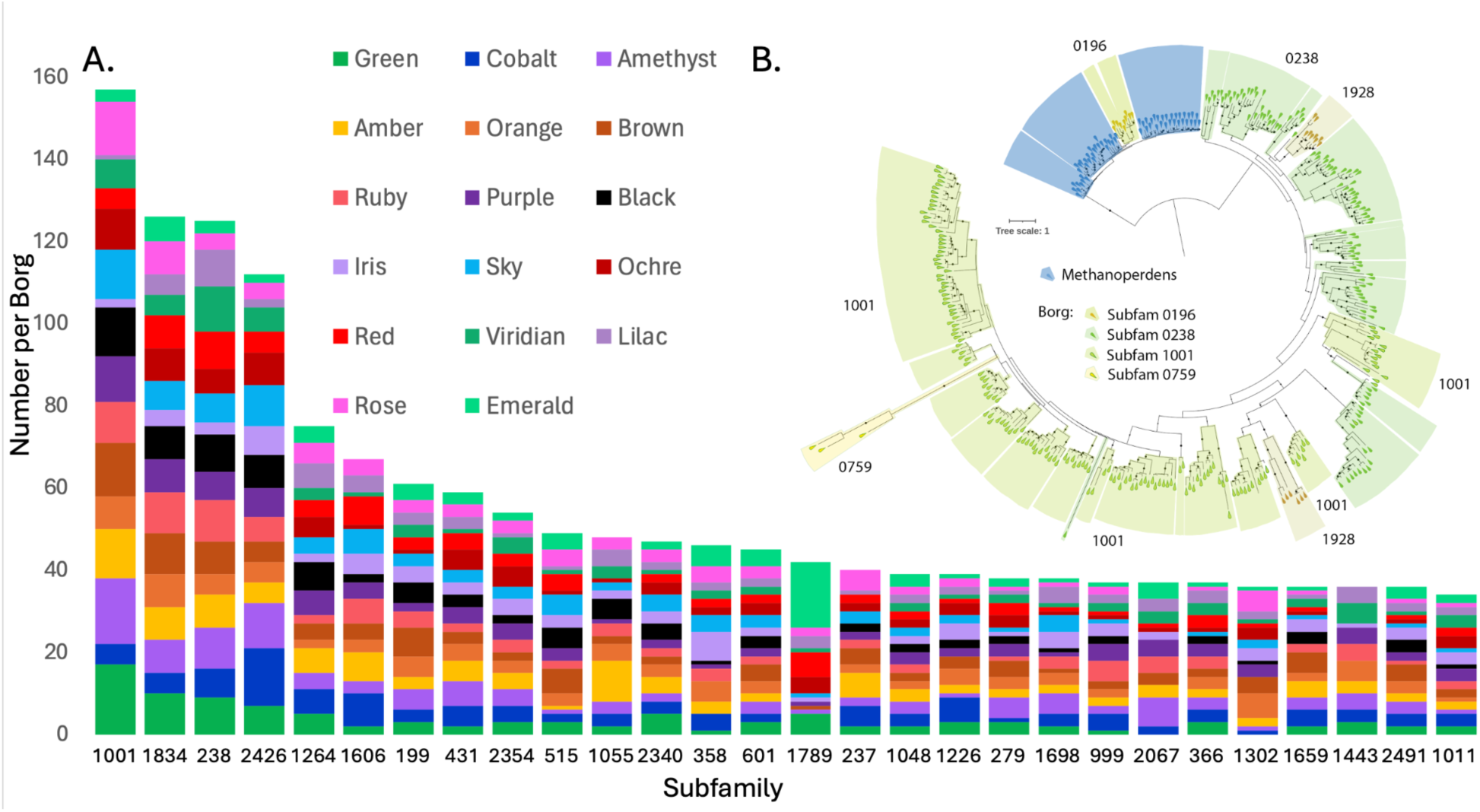
A. Number of proteins assigned to each highly multicopy protein subfamily for each of the 17 Borg genomes based on clustering reported by (Schoelmerich et al. 2024). Each distinct Borg has a color-based name. **B**. Maximum-likelihood phylogenetic tree showing intermixing of Borg proteins from four of these subfamilies, supporting their treatment as a single group (Group 1). A few sequences without subfamily clusters were assigned to subfamilies based on phylogeny. Notably, subfamily 196 places within a large clade of *Methanoperedens* sequences, suggesting that Borgs acquired these 10 sequences by recent lateral transfer from *Methanoperedens*. Representatives of all subfamilies have the expected structures and active site residues (including 0759). A few proteins were too poorly folded to enable confident analysis, but in all cases, the protein sequences phylogenetically placed in clusters of proteins with the expected structures.

Group 1 proteins share a common fold with eukaryotic JAMM (AMSH) deubiquitinases (**Figure 2A**). These Zn-metalloproteases liberate ubiquitin from target proteins using a highly conserved catalytic core comprising a nucleophilic Ser and metal ion coordinating His/His/Asp or Glu residues (Suresh et al. 2020); **Supplementary Information**). Although deubiquitinases are known to remove ubiquitin from a variety of biomolecules, predicted cytoplasmic localization suggests that the targets are proteins. We mapped sequence conservation from the alignment of 315 Borg sequences onto the predicted structures and found that conservation localized to the expected active site, which is adjacent to the beta sheet face (**Figure 2A**). Most of the 315 Borg proteins are likely metalloproteases with the expected active site residues (e.g., **Figure S3A**). The few with active site variants (e.g., Amber 1107 and Amber 1231, **Figures S3B, 3C**) may have modified functionality.

**Figure 2:**
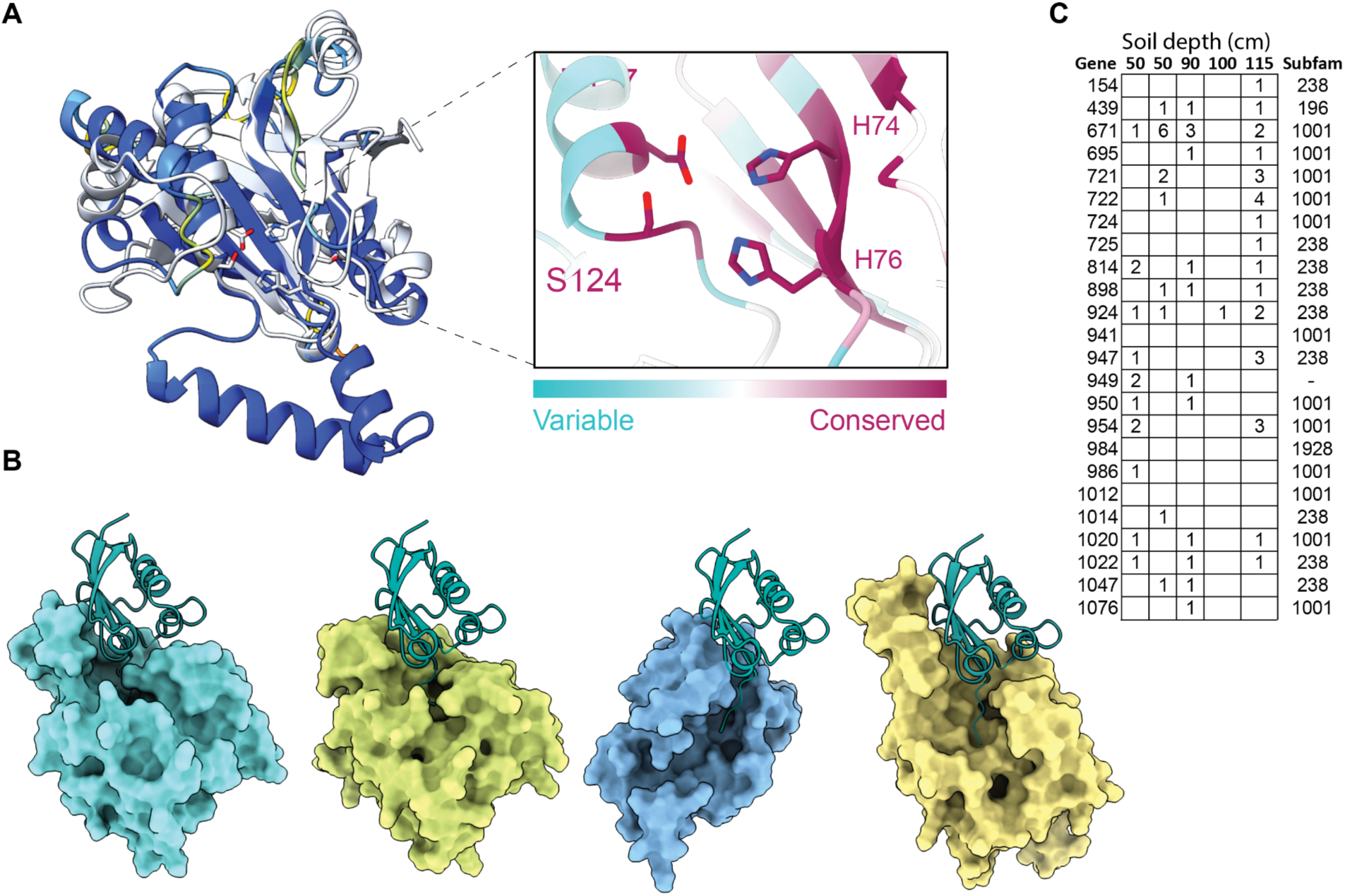
Putative de-SAMPylation enzymes. **A**. Human AMSH deubiquitinase PDB 2znr) and Borg deSAMPylase (Orange 866) colored by AlphaFold2 pLDDT confidence (blue is high confidence). The inset shows the conserved Borg protein active site residues expected for Zn metalloprotease and is colored by sequence conservation, as defined using an alignment of Borg putative de-SAMPylation enzymes. **B.** AlphaFold3 predictions for Orange Borg deSAMPylase proteins (surface representation from left to right: 883, 866, 892, 885) bound to *Methanoperedens* SAMP (teal). **C.** Metatranscriptomic read counts (from a published nanopore dataset; (Schoelmerich et al. 2024)) for Black Borg de-SAMPylases in five soil samples and the subfamily affiliations of each protein. Expression was detected for up to 21 of the 24 proteins. The gene numbers reveal that these multicopy proteins are often encoded in close proximity.

Unlike Eukaryotes, Archaea generally do not use ubiquitin but instead post-translationally label their proteins for proteolysis using a Small Archaeal Modifier Protein (SAMP; (Darwin and Hofmann 2010). Using the sequence of the characterized *Haloferax* SAMP (Humbard et al. 2010), we identified SAMP with the expected conserved ꞵ-grasp fold and di-glycine tail encoded in *Methanoperedens* genomes (e.g., cMp 2846). An E1 SAMP-adding enzyme is encoded in the cMp genome (3305) and co-folding predicts it to bind the cMp SAMP into the active site (**Figure S4)**. Borg deSAMPylase-like proteins also bind the cMp SAMP and also position the di-glycine tail into the active site (**Figure S4B**). Taken together, we suggest that Group 1 Borg proteins are deSAMPylases.

The complete ∼ 4 Mbp *Methanoperedens* genome (cMP) has two putative deSAMPylases with the expected topology and conserved active site residues. cMP 1357 falls within a clade defined by sequences from *Methanoperedens* and other archaea whereas cMP 650 falls within a clade of other *Methanoperedens* sequences that includes a subclade comprised of 10 Borg proteins (mostly subfam0196; **Figure 1A**). The most likely interpretation is that Borgs acquired these 10 deSAMPylases from *Methanoperedens* via (recent) lateral transfer. This reinforces the established pattern involving sharing of proteins between Borgs and *Methanoperedens (Al-Shayeb et al. 2022),* supporting their physical association.

An obvious question is why Borg genomes encode so many deSAMPylases. There is substantial predicted variation in the shape of the region that would bind the SAMPylated protein (**Figure 2B**), so distinct variants may deSAMPylate different targets. Specificity may be enhanced by structural rigidity conferred by disulfide bonds distant from the active site that are predicted in almost all examples (often three or four cysteine pairs per protein). Supporting complementary functionality, available nanopore transcript data for Black Borg (Schoelmerich et al. 2024) show that multiple variants were expressed *in situ* (**Figure 2C**). Recent results indicate that bacteria add ubiquitin to proteins to block phage packaging and interfere with reinfection (Hör et al. 2024), so deSAMPylation may be an analogous response to an archaeal host SAMP-based defense mechanism.

Given the proliferation of Borg deSAMPylases and long branch lengths in the tree suggestive of rapid divergence, we suspected that these multicopy genes are under increased diversifying selection relative to those of *Methanoperedens*. The dN/dS ratio for the large Borg clade (**Figure 1B**) is 0.235, compared to 0.195 for *Methanoperedens* and 0.172 for the Borg clade nested within the *Methanoperedens* group. Although none of these values is indicative of positive selection (dN/dS > 1), use of dN/dS as a purely comparative metric suggests that selection is relaxed in genes of the large Borg clade. Using the RELAX function in Hyphy (Kosakovsky Pond et al. 2020), we calculated a relaxation parameter, K, of 0.53, which was verified to fit the observed data with a p-value of 0.0000**. Therefore, we conclude that relaxation of negative selection is significant in the large Borg clade (K < 1, p < 0.05).

### Borg investments in glycoconjugates

Structural prediction revealed many multicopy genes with functions likely related to surface modification, particularly production of glycoconjugates. Roles in N-glycan biosynthesis are suggested for diverse members of subfam0199 (**Figure 3A)**, whose predicted structures correspond well with those of dolichyl phosphate mannose synthase (DPMS; **Figure 3B)**. This protein catalyzes transfer of mannose from GDP-mannose to the dolichol carrier Dol-*P* (an isoprenoid) to yield dolichylphosphate mannose, likely for decoration of external proteins. Surface representation of the Borg proteins reveals a volume that could accommodate the UDP-mannose donor substrate with the donor mannose group positioned proximal to the lipid entry channel (**Figure S5A,B**). Nanopore metatranscriptomic data (Schoelmerich et al. 2024) demonstrate expression of four of the five Black Borg DPMS proteins in wetland soil, consistent with complementary functionality.

**Figure 3:**
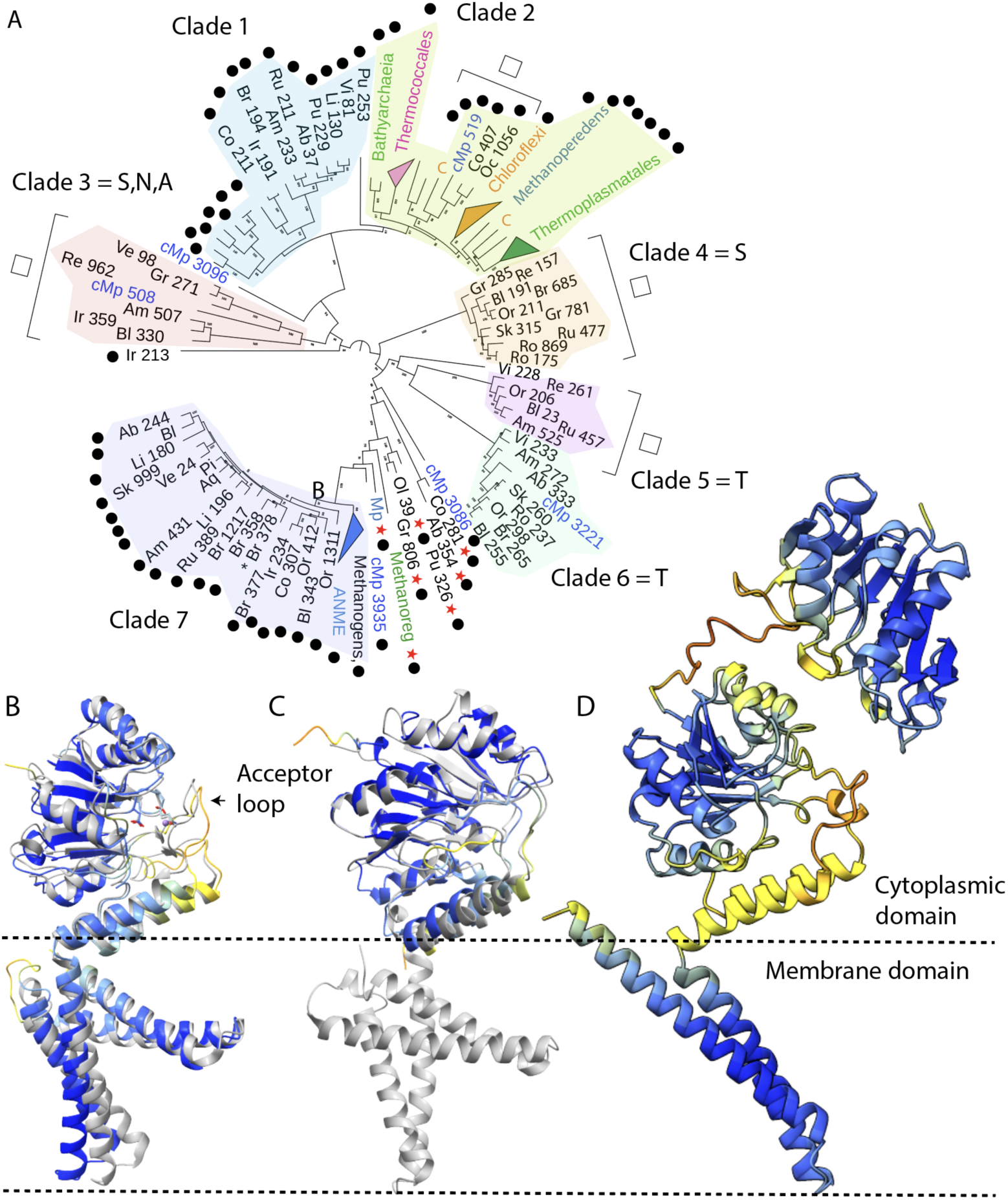
Borgs encode multicopy proteins related to dolichyl phosphate mannose synthase, implicated in glycosylation. **A**. Phylogenetic tree of putative DPMS proteins (first two letters indicate the Borg name (except for Amber, Ab), followed by the gene number). Almost all unnamed sequences are from archaea. C = Chloroflexi, cMp=complete *Methanoperedens* genome. Black dots indicate DPMS with metal-binding residues. Residue(s) in place of glutamine are indicated in the clade name. Open boxes indicate groups that lack the membrane anchor and red stars identify proteins with extra cytoplasmic domains. The asterisk indicates a split protein likely functional following +1 frameshift. Some proteins may have been transferred to Borgs from *Methanoperedens* (e,g, Clade 7) and others from Borgs back to the *Methanoperedens* (e.g., cMp in Clade 6). **B.** Cobalt 211 aligns very well to PDB 5mm0, and has all 40 conserved residues identified of the reference structures. **C**. Some putative DPMS, e.g., Orange 206, lack the membrane anchor, but align well in the cytoplasmic region. **D.** Other putative DPMS, e.g., Green 806, have an extra cytoplasmic domain. In **B,C**, PDB 5mm0 is gray and in **B,C,D,** the Borg proteins are colored by the AlphaFold plDDT scheme (darker blue indicates higher confidence).

The DxDxQ (particularly DxQ) residues of the metal binding motif required for DPMS functionality (Gandini et al. 2017) occur in many, but not all, phylogenetically defined Borg clades, and in the DPMS from *Methanoperedens* and other archaea (**Figure 3A; Supplementary Information**). There is variability in the number of alpha helices in the membrane anchor, and 13 of the 17 Borgs have at least one DPMS that completely lacks transmembrane alpha helices (**Figure 3C**). Only members of one clade that lack the alpha helical anchor have the DxDxQ motif (**Figure 3A**). Like all DPMS, anchorless variants have amphipathic helices with a hydrophobic face (**Figure S5C)** that likely enables localization at the cell membrane. Some anchorless variants are likely functional if they have the conserved DxDxQ motif, given the functionality of truncated DPMS from *Pyrococcus furiosus (Gandini et al. 2017)*. However, the active site residues are different in some anchorless variants (**Figure S5E**).

Adding to DPMS diversity, five Borg and two archaeal proteins have an extra domain that aligns to the primary cytoplasmic domain of DPMS (**Figure 3D**, **Figure S6A,B).** Alphafold sometimes predicted homo-tertrameric structures (**Figure S6C**, albeit with modest confidence scores), recreating the known tetrameric eukaryotic homolog structure (Ardiccioni et al. 2016). All seven proteins with an additional cytoplasmic subunit have metal binding sites in the main cytoplasmic subunit and two also feature the metal-binding DxDxQ motif in the extra domain (**Figure S6D**). One predicted protein with the DxDxQ motif is split (Brown 377,378), but it should be functional following a +1 frameshift (**Figure S7**). Overall, the structural diversity hints at broad functionality that may be achieved via proteins with and without a membrane anchor, via a second cytoplasmic domain, and via multimerization as homo- or heteromers.

Consistent with the pattern for deSAMPylases, the Clade 7 Borg DPMS phylogeny (**Figure 3A**) is suggestive of diversification following acquisition of Borg proteins via lateral gene transfer from *Methanoperedens* or related archaea. However, single *Methanoperedens* DPMS-like proteins within Clades 3 (6 Borg proteins) and 5+6 (14 Borg proteins) suggest that the reverse process, in which DPMS were acquired by *Methanoperedens* from Borgs, also likely occurs. The dN/dS values for Borg proteins = 0.121 compared to 0.098 for *Methanoperedens*, with a relaxation parameter, K, of 0.76, and p = 0.0241**, consistent with significant relaxation of negative selection in Borgs (K < 1, p < 0.05). Thus, it now appears that variants that undergo diversification in Borgs can make their way back into host genomes.

Two DPMS genes occur in Cobalt Borg in a region that is enriched in genes seemingly involved in cell surface decoration (**Figure 4**). Similar regions occur in all Borgs. The Cobalt Borg region encodes multiple UDP-sugar-epimerases, glycosyltransferase MshA, deacetylases, and genes that may be involved in biosynthesis of mycothiol, a complex N-acetylglucosamine-based polymer. Also present is a gene cluster for which structural analysis supports a role in biosynthesis of the polysaccharide O-methyl phosphoramidate, which contains unusual C-P bonds, genes that may form phosphonates, and others with predicted roles in phospholipid and glycerophospholipid metabolism (**Supplementary Information**). Eight of the forty Cobalt Borg proteins analyzed (**Figure 4**) include SEC signal sequences (for localization and processing at the host membrane) and eight are predicted to be membrane-embedded. These findings suggest that Borgs invest in production of diverse and unusual glycoconjugates, likely for surface decoration.

**Figure 4:**
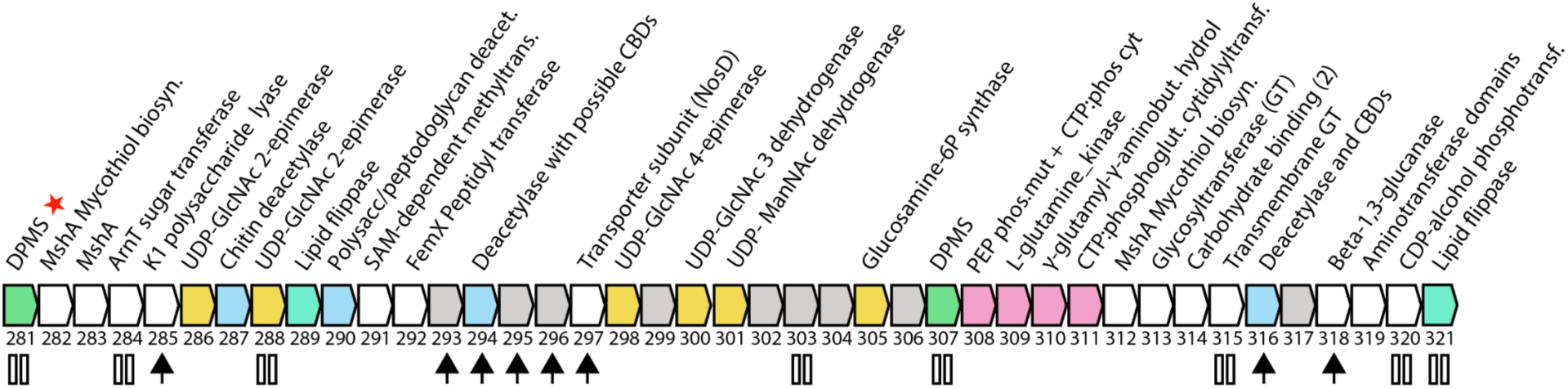
Borg genomes encode regions enriched in proteins for production of glycoconjugates and other cell surface polymers. A region of the Cobalt Borg genes (genes 281-321) encodes genes for N-glycan production (green, aqua; red star indicates that the DPMS has an extra cytoplasmic domain), with functions related to metabolism of UDP-N-acetylglucosamine (GlcNAc) and similar compounds (yellow), as well as deacetylases (blue) and proteins related production of O-methyl phosphoramidate (pink) (e.g., 311: CTP:phosphoglutamine cytidylyltransferase) and phosphonates (308 is likely a fusion of phosphoenolpyruvate mutase and CTP:phosphoglutamine cytidylyltransferase). CBD carbohydrate binding domain. Paired vertical lines indicate transmembrane domains and arrows denote a SEC signal sequence.

### Investment in encapsulation

Multicopy subfam2354 and subfam1011 (**Figure 1**) are profiled as viral capsid proteins, paralleling a prior report of Borg proteins similar to those of an archaeal virus (Schoelmerich et al. 2024). The capsid-like proteins are often most similar to those of *Haloarcula californiae* HCIV-1, an archaeal icosahedral virus with a linear genome, an internal membrane and a tailless icosahedral morphotype that includes hexamers of a single jelly roll (SJR) protein. In all 17 Borgs, a syntenous region encodes one capsid-like protein and four or five hypothetical proteins. For Black Borg, the only Borg for which extensive transcript data were recovered from an Illumina dataset generated for an 80 cm depth soil sample, the SJR protein is 801. 801, 799 and 800 are exceedingly highly expressed; in fact, 76% of the Black Borg transcripts map to this locus (allowing 2% SNPs, **Figure 5A**). The analogous Purple Borg genes are also relatively highly expressed. Similar caspid-bearing gene clusters also occur in some mini-Borgs (Borg-like ECEs with genomes ∼10x smaller than Borgs (Shi et al. 2024) **(Figure S8**), and some are expressed. Major capsid proteins are typically among the most highly expressed genes in viruses (Moniruzzaman et al. 2017), supporting the inference that these proteins are involved in encapsulation.

**Figure 5:**
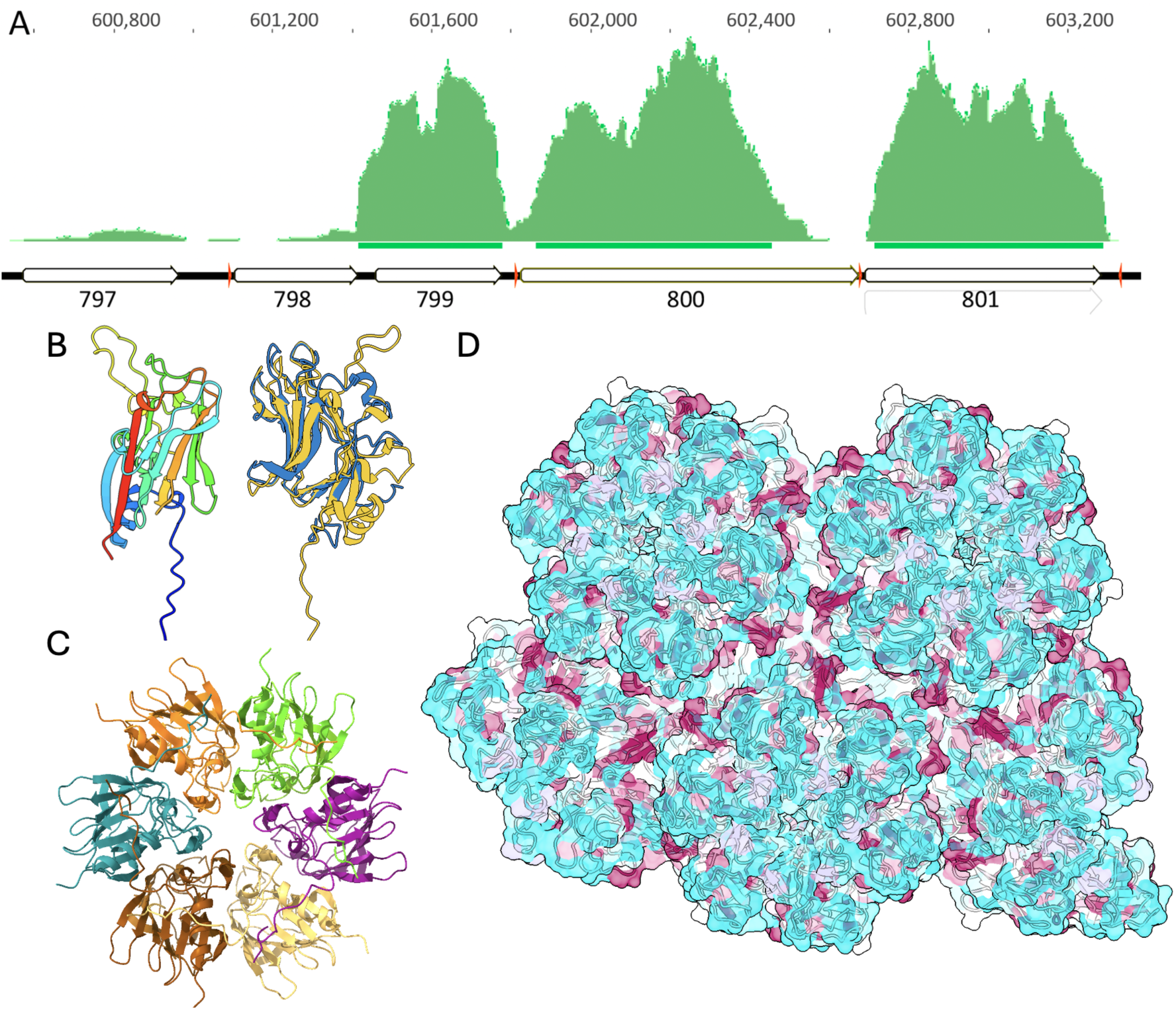
Black Borg has a very highly expressed capsid-like protein that multimerizes into capsid-like arrays. **A.** Metatranscriptomic reads mapped to the Black Borg genome primarily localize to three genes, the third of which is a putative capsid protein (801). Red tick marks are ribosome binding sites. **B**. Black 801 monomer colored by position (start blue, end red) and, in gold, aligned to the reference 6h9c VP7 subunit M. **C**. Hexamer prediction for Black 801 colored by sequence conservation (red is high conservation; ipTM = 0.8, pTM = 0.82). **D.** One outcome for a prediction of the 24-mer of Black 801 showing assembly of hexamers into a capsid-like sheet, with sequence conservation (red) at the hexamer junctions.

The putative capsid protein Black 801 (**Figure 5B)** is predicted to assemble into hexamers with high confidence (**Figure 5C)**. Scores for multimers with fewer subunits are much lower, indicating that hexameric assembly is probable. In some simulations, rings composed of five or six monomers form extended sheet-like arrays, with conserved amino acids at interfaces between hexameric units, as expected for capsids (**Figure 5D**). The inconsistency in multimer conformations, which we also demonstrated for HCIV-1 major capsid proteins, aligns with the known limitations of AlphaFold2 and AlphaFold3 for predicting higher order assemblies (Bryant et al. 2022). Overall, the extremely high level of expression and consistent hexameric assembly (as known for HCIV-1) align with capsid-like behavior.

In addition to the region encoding 810, Black Borg also encodes five consecutive SJR-like proteins (867 - 871) and proteins with similar folds occur sequentially in 12 other Borg genomes (**Table S9**). For example, Amethyst Borg encodes six consecutive SJR fold proteins (1053 - 1058). We predicted a variety of multimers of all six Amethyst Borg proteins but only found near-perfect scores for hexamers (**Figure S9**). These findings indicate a surprising variety of potential capsid-like proteins in Borg genomes.

Some mini-Borgs also encode an additional SJR-like protein, and these define a large phylogenetic cluster that includes one Black Borg protein from the cluster of capsid-like genes (867). Overall, 867 shares 69 - 80% amino acid identity with 20 different proteins from two different mini-Borg types, yet only 39% identity with the homolog from the most closely related Borg (Ochre). This provides evidence for recent lateral transfer of capsid-like proteins between Borgs and mini-Borgs.

Eight Borgs have an additional gene cluster that typically encodes three proteins, two of which are nearly isostructural, and two genomes have duplicates of this region in close proximity (**Table S9**). Structural features and distant homology identified by structure-based HMM model refinement (see Methods) suggest tail- or capsid-like functions. This gene cluster and the two previously discussed gene clusters are typically encoded within an ∼100 gene interval (**Table S9**, suggesting consistent co-localization of functions likely related to encapsulation.

Other multicopy proteins encoded in all Borg genomes (subfam1773) resemble a bacteriophage HS1 tail needle knob protein (4k6b), but are more likely to be capsid-related jelly roll fold, membrane anchored proteins (**Figure S10**). Like 4k6b, they assemble into homo-trimers with moderate confidence (**Figure S10**), and other multimers with much lower confidence. The Borg proteins display a hydrophobic alpha helical region that may insert into a membrane.

### Borg characteristics resemble those of giant eukaryotic viruses

Credible capsid-like proteins, the presence of viral-like (Herpesvirus) replication machinery (Perkins et al. 2008), combined with linear genomes of large size motivated a comparison of Borgs and giant eukaryotic viruses. Giant eukaryotic viruses are now referred to as members of the phylum *Nucleocytoviricota (Chihara et al. 2022)*, but were previously described simply as nucleo-cytoplasmic large DNA viruses (NCLDV; (Iyer et al. 2006); (Rigou et al. 2024). Like Borg genomes, many *Nucleocytoviricota* genomes are terminated by long inverted repeats, yet their coding structure is different, as genes frequently alternate between strands. Similar to Borgs, many larger *Nucleocytoviricota* (e.g., Mimivirus and Megavirus) have lower %G+C content compared to their hosts (Wilhelm et al. 2017). *Nucleocytoviricota* exhibit high levels of horizontal gene transfer with other viruses and organisms, a Borg feature that led to their naming. These acquired genes often come from the hosts, in the case of Borgs from *Methanoperedens* archaea.

Arguably the most striking feature of Borgs is their large inventory of genes that are normally only associated with organisms (Al-Shayeb et al. 2022; Schoelmerich et al. 2024), **Table 1, Table S10**), also true of giant eukaryotic viruses (Moniruzzaman et al. 2023). As in *Nucleocytoviricota (Mönttinen et al. 2021)*, Borg inventories of sugar and lipid-related genes may be involved in decoration of capsid-like structures. In fact, given this, and their diverse inventory of putative capsid-like proteins, Borgs may construct capsids that are analogous to the complex structures reported recently for *Marseilleviridae*, which have eight protein components and an internal membrane, likely with a glycoprotein surface (Moniruzzaman et al. 2023). As for *Nucleocytoviricota*, Borgs encode genes for coenzymes, and nine Borgs have three sequential genes for production of NAD. NAD production may counter host defense mechanisms that deplete cellular NAD levels (Osterman et al. 2024). Borgs and *Nucleocytoviricota* both encode genes for glycolysis / gluconeogenesis (in Borgs, up to seven genes, including for sequential steps encoded by sequential genes) and related genes of the pentose phosphate pathway. Other Borg proteins are involved in pyruvate, phosphoenolpyruvate, and acetyl-CoA metabolism, and the TCA cycle (e.g., sequential citrate synthase and aconitase genes, **Table S10**, also see (Schoelmerich et al. 2024). Thus, overall, investments in core carbon metabolic processes are features of both Borg and large *Nucleocytoviricota* genomes (Moniruzzaman et al. 2020).

**Table 1.**
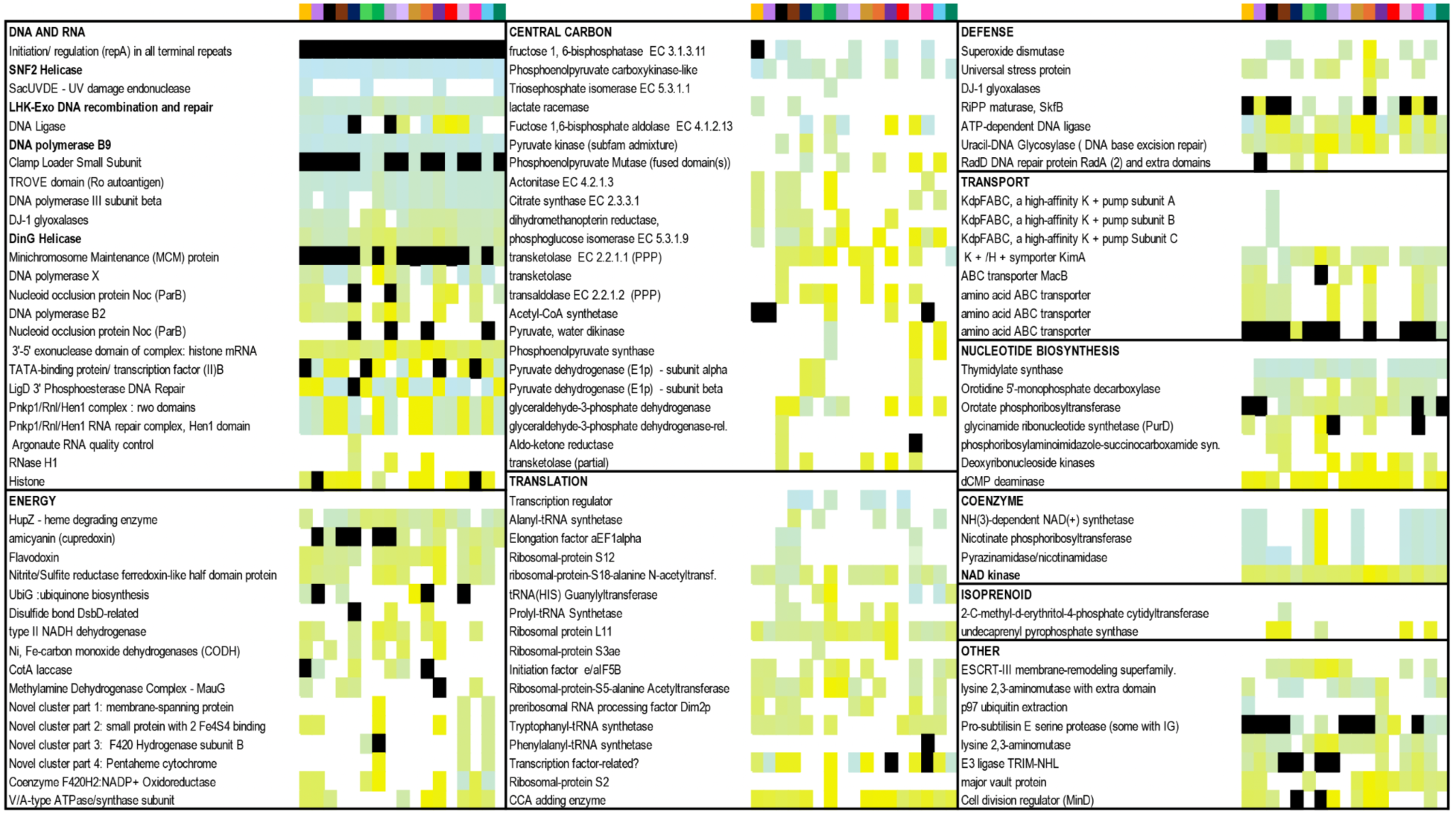
Predicted protein functions across the 17 Borg genomes (columns ordered alphabetically), excluding the multicopy genes described elsewhere in the text and functions discussed in detail previously (e.g., multiheme cytochromes). Rows are gene names and genes are grouped by function type. Genes in most of these categories occur in giant eukaryotic viruses. Genes that occur in multicopy in a genome are indicated in black (positions not indicated). Within each functional group, genes were sorted based on average position (approximately so as to not break up sequential genes). Color indicates presence and the shade of color indicates gene position in the genome, from early (blue) to late (gold); for details see **Table S10**. Although there is variation, genes for the same function tend to occur in similar genomic regions.

Unlike most *Nucleocytoviricota*, Borgs do not encode RNA polymerases, but they do encode many transcriptional factors to recruit host machinery to drive transcription. Both *Nucleocytoviricota (Brandes and Linial 2019; Raoult et al. 2004)* and Borgs possess extensive translational machinery. For example, Borg genomes encode up to 23 tRNAs, 14 have one to three tRNA synthetases, they have one or two of four different ribosomal proteins and 15 encode a CCA-adding enzyme. Interestingly, similar genetic repertoires occur in some giant viruses of bacteria (megaphages; (Al-Shayeb et al. 2020). Like some *Nucleocytoviricota*, Borgs have genes for many other DNA and RNA functions, including for nucleotide biosynthesis. Similar to *Nucleocytoviricota*, they have genes for transport of K+ (Moniruzzaman et al. 2023) and other compounds (Aylward et al. 2021). Also paralleling *Nucleocytoviricota (Brandes and Linial 2019)*, some Borgs have genes for radical defense (e.g. superoxide dismutase) and nucleic acid damage repair.

*Nucleocytoviricota* genomes typically encode DNA polymerase B (Aylward et al. 2021). All Borgs encode at least one DNA polymerase B, but sequences do not place with those of Eukaryotic viruses, so a common ancestor of Borgs and *Nucleocytoviricota* is not phylogenetically supported (**Figure S11**). Consistent with this distinction, Borg capsid-like proteins are most comparable with those of archaeal *Sphaerolipoviruses*, not the capsids of *Nucleocytoviricota*.

*Nucleocytoviricota* typically encode proteins to dynamically package DNA into chromatin (Liu et al. 2021; Gaïa et al. 2023). Many Borgs have one or more histone proteins, histone remodeling helicases, histone protein methyl/acetyl transferases, and histone protein demethyl/deacetyl transferases. Further underscoring an investment in high-order DNA compaction, Borgs all have a gene encoding 3’hExo ternary complex, an exonuclease that negatively regulates the abundance of histone mRNA to appropriately titrate histone concentrations during DNA replication (Dominski et al. 2003). Interestingly, histone genes sometimes co-occur with a gene for a predicted vault protein (the subunits of which are predicted to assemble into vault-like arrangements). Less confidently functionally annotated are almost ubiquitous genes for a tubulin / FtsZ-like protein (**Supplementary Information**). Tubulin-like proteins occur in some *Nucleocytoviricota* (Da Cunha et al. 2022) and in some bacteriophages, tubulin (PhuZ)-based spindles move the capsids to the surface of a phage nucleus for genome packaging (Chaikeeratisak et al. 2021). Packaging ATPases (similar to those of Phycodnaviridae, (Mönttinen et al. 2021) could only be confidently identified in some mini-Borg genomes and one partial Apricot Borg genome. There are multiple ATPase-like proteins in Borg genomes, some with structures resembling packaging ATPases, but their predicted functions vary. Thus, the presence of packaging machinery in Borg genomes remains unresolved.

Like *Nucleocytoviricota (Filée 2009)*, Borg genomes are peppered with genes for selfish genetic elements (e.g., transposons). In fact, genes for TnpB, IsrB and Cas12 are among the most prevalent multicopy proteins in Borg genomes (**Figure S12**).

Ubiquitous, tandemly repeated nucleotide sequences are striking features of both Borg genomes and genomes of a diverse group of *Nucleocytoviricota* (**Figure 6**). Such repeats are very rare in the curated and complete *Methanoperedens* genomes, and occur ∼ 3 to ∼5 times less frequently (as microsatellites) in some bacterial genomes (Subirana and Messeguer 2020). The tandem repeat patterns in *Nucleocytoviricota* and Borgs are similar in terms of their distributions, the number of repeats per region, and average repeat length (**Table 2, Table S11 - S14**). All but the shortest repeats are almost always novel to each region, and population-level variation in unit repeat number per locus in both is indicative of rapid evolution.

**Figure 6:**
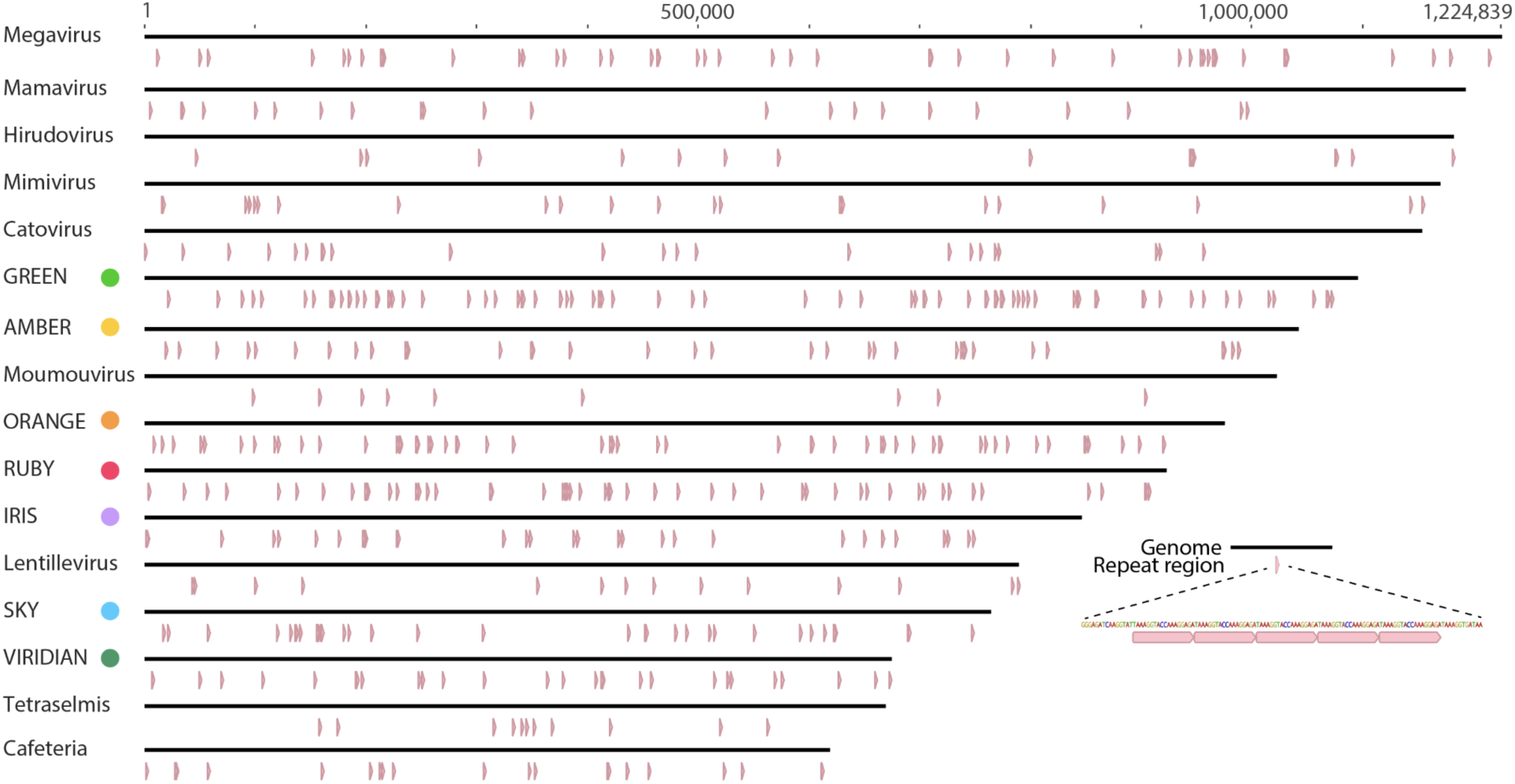
Borgs and *Nucleocytoviricota* display generally similar abundances and distributions of perfect tandem repeats. Genomes for examples of different types of *Nucleocytoviricota* and a selection of Borgs (colored dots), listed in order of decreasing genome length. For more complete information see **Tables S11-S14**.

**Table 2:**
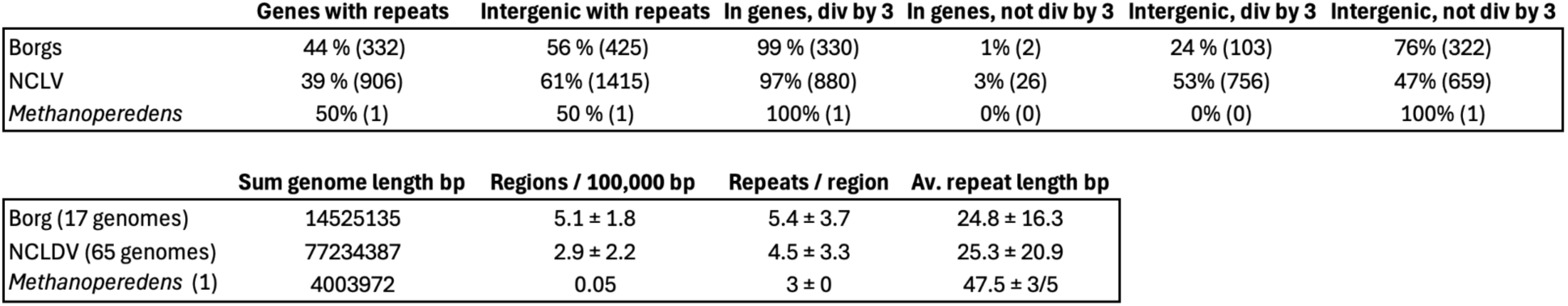
Overview of tandem repeats in Borgs, *Nucleocytoviricota* and the complete *Methanoperedens* (cMp) genome. Tandem repeat regions were inventoried if they were at least 50 bp long with at least 3 repeat repeat units and a minimum unit repeat length of 9 bp.

Of the tandem repeats, 44% and 39% occur within genes in Borgs and *Nucleocytoviricota*, respectively, and they almost always generate amino acid repeats (**Table 2**). In Borg genomes, amino acid repeats often introduce intrinsic disorder (Schoelmerich et al. 2023). This is also suggested for some *Nucleocytoviricota* repeat proteins, but most are predicted to form ankyrin repeats or other alpha-solenoid structures (Mönttinen et al. 2021). Importantly, it is not uncommon for repetitive regions within proteins to be predicted as both intrinsically disordered and alpha-helical (Mier et al. 2020). In ankyrin, the alpha solenoid structure is an emergent property of multiple intrinsically disordered repeats interacting with each other. In other cases, multimerization or post-translational modification cause low complexity regions to toggle between disordered and alpha-helical structures (Kumar and Balbach 2021).

The ability to transition between disordered and ordered states is characteristic of proteins that undergo liquid-liquid phase separation to form separate cellular compartments that are not membrane bound. Recent work has claimed that the viral factories (Rigou et al. 2024) of the *Nucleocytoviricota* Mimivirus and Noumeavirus are phase separated organelles composed of, and scaffolded by, intrinsically disordered proteins (Aftab et al. 2024). Sequestration of viral replication also occurs through the production of a ‘phage nucleus’ in some jumbophage (Prichard et al. 2024), possibly suggesting similar adaptations in large viruses. The high number of intrinsically disordered proteins in Borgs may hint at their ability to form phase-separated compartments similar to the *Nucleocytoviricota* viral factories.

### Clues to Borg evolution

Borgs genomes feature many multicopy genes. In the case of putative deSAMPylases, genomic proximity suggests origination by gene duplication (**Figure 2C**). Other multicopy functions relate to protein decoration, nucleic acid manipulation, encapsulation and redox activities (**Figure S12**). Multicopy genes also make up a large proportion of *Nucleocytoviricota* gene content (Mönttinen et al. 2021). In fact, protein duplication is a suggested driver of viral gigantism (Brandes and Linial 2019; Machado et al. 2023). In *Nucleocytoviricota*, the “genomic accordion” process is suggested to maintain or expand host range (Van Etten 2003; Raoult et al. 2004; Elde et al. 2012). Should the evolutionary model of giant size through gene accretion extend to Borgs, mini-Borgs (with their ∼10X smaller genomes, (Shi et al. 2024) may resemble an ancestral state.

We noted consistent genomic locations for many genes that occur in two or more Borg genomes and are single copy per Borg genome (1931 examples). For example, DNA polymerase B9 family genes are all encoded early (0.08 ± 0.2 along the genome) and the widespread CCA-adding enzymes are consistently encoded ∼75% of the way along the genome (0.73 ± 0.11). In fact, 64% of the 1931 genes our at the same average position with a standard deviation of ≤ 0.1 (**Figure S13**), and 251 occur in such close proximity in 7 or more Borgs (**Table 1, Table S10)**. This greatly extends the previously noted pattern of 40 universally present single copy Borgs genes in generally similar genomic positions (Schoelmerich et al. 2024). Often, but certainly not always, the phylogeny of genes in similar genomic locations matches the phylogeny based on the 40 universal single copy proteins. The findings indicate extensive inheritance of a genetic backbone, shaped by gene loss.

The presence of many of the currently existing Borg genes in a common Borg ancestor may align with the speculation that the functional gene-rich Borg backbone was acquired from an archaeon that shared a common ancestor with *Methanoperedens* (Schoelmerich et al. 2024)). It has been similarly suggested that giant eukaryotic *Nucleocytoviricota* arose via gene loss from an organismal source (Brandes and Linial 2019), but most phylogenetic evidence tends to support the evolution of viral gigantism from smaller viruses (Koonin and Yutin 2019; Iyer et al. 2006; Moreira and Brochier-Armanet 2008). Alternatively, it is possible that the current pattern of gene position conservation and sequence reflects inheritance from a gene-rich ancestral ECE, possibly one that evolved at the time of origination of the *Methanoperedens* genus, with subsequent lineage-specific gene gain and loss (Iyer et al. 2006; Claverie and Abergel 2013). Regardless of the details, it appears that the last common Borg ancestor had a giant genome.

So, are Borgs giant archaeal viruses? They are undoubtedly archaeal, which makes the question: are Borgs viruses? Periodic viral-like bursts are suggested based on very high, and highly variable, Borg to host chromosomal DNA copy numbers in some samples. These phenomena may also reflect selective preservation of encapsulated Borg DNA (Schoelmerich et al. 2024). The small number of genes encoded within the known archaeal viruses precludes a detailed comparison with Borg gene content and the lack of a ubiquitous packaging ATPase and for evidence of full capsid assembly limits the argument for classification of Borgs as viruses. However, Borgs encode functions that are similar to those of viruses in general, such as a SNF family helicase (possibly involved in chromatin remodeling), proliferating cell nuclear antigen (sliding clamp DNA polymerase), and DNA polymerase B, and the genomic placements are generally analogous. The DNA polymerase B genes are encoded very early in the large replichore whereas genes involved in translation and ∼100 kbp regions that encode many putative structural genes occur mid-genome, and histones that are probably required for organization and packaging of DNA (**Figure S14**) are encoded towards the end of the large replichore or within the small replichore (**Table S10**). ESCRT-III-like proteins are present in some Borgs (**Figure S15**). These are involved in membrane remodeling, and in *Nucleocytoviricota* generate the lipid component of capsids.

Overall, Borgs have numerous features suggestive of lifestyles somewhat analogous to those of giant eukaryotic viruses and we conclude that the weight of evidence now supports a more viral-like than plasmid-like lifestyle. Regardless of the classification of Borgs, the parallels with *Nucleocytoviricota* are striking. Given no phylogenetic evidence for common ancestry, many *Nucleocytoviricota* and Borgs similarities are likely due to convergent evolution. Gigantism in eukaryotic viruses evolved independently in separate lineages (Koonin and Yutin 2018, 2019), likely occurred repeatedly in bacteriophages to generate the many distinct clades (Al-Shayeb et al. 2020), and may have occurred again in virus-like Borgs of archaea.

Koonin et al. write “In a sense, all evolution of life is a history of coevolution between MGEs and their cellular hosts” (Koonin et al. 2020). Several studies have suggested that ancient eukaryote-virus gene exchange shaped the early evolution of complex cells, possibly even leading to traits such as linear chromosomes or the nucleus (Takemura et al. 2015; Villarreal and DeFilippis 2000). Direct evidence for early virus-to-eukaryote gene exchange has remained elusive and an archaeal virus that could serve as a clear progenitor in these evolutionary scenarios has not been found. It is intriguing to consider that archaeal evolution at around the time of eukaryogenesis may have been shaped by large and complex viruses. Given that Borgs lack clear phylogenetic affinity for eukaryotic viruses and eukaryotic replisome components, it is unlikely that they represent the missing link in early eukaryotic evolution. However, Saturn mini-borgs encode a delta-like family B DNA polymerase that has phylogenetic placement proximal to eukaryotic polymerases alpha, zeta, and delta (**Figure S11**). This raises the possibility that these elements may have played a role in the early emergence of the eukaryotic replisome. Regardless, the existence of Borgs and mini-borgs opens the door to future work on other lineages of mobile elements and complex archaeal viruses that may have played a role in the emergence of cellular complexity.

## Methods

### Identification and analysis of multicopy subfamily proteins

Multicopy subfamily analysis was conducted by analysis of the subfamilies reported for the 17 complete and highly curated, near-complete Borg genomes that we reported, see ((Schoelmerich et al. 2024)). Subfamilies for which functional and structural predictions were similar or the same (see below) were evaluated using phylogenetic analysis, and categories grouped as appropriate. Analyses did not rely upon strict subfamily assignments because structural alignments demonstrated similar folds for sequences from different subfamilies. (and *vice versa*).

### Protein structure prediction and functional analysis

The genomes of Orange, Black, Green, Amber, Amethyst, Cobalt and Ruby Borgs were selected as representative of Borg diversity based on their relatedness, as inferred from the phylogeny based on 40 single-copy universal genes that we reported previously (Schoelmerich et al. 2024). For each protein in these seven Borg genomes, ColabFold (--amber --templates --num-recycle 3) (Mirdita et al. 2022) run locally was used to predict AlphaFold2 structures that were evaluated for similarity to proteins in the Protein Database (PDB, https://www.rcsb.org/). Multimer calculations were performed using either AlphaFold2 multimer (Evans et al. 2022) and AlphaFold3 (Abramson et al. 2024). Output models were assessed by pLDDT (>70 overall model pLDDT considered confident), PTM (>0.6 considered confident), and iPTMscores (>0.8 considered confident). We prioritized analysis of structures for proteins in prevalent multicopy subfamilies. In all instances, predicted structures were searched against the Protein Data Bank (PDB) using Foldseek (https://search.foldseek.com/search). Specific protein structures of interest were additionally searched against the online Foldseek database (https://search.foldseek.com/search). Predicted structures were visualized primarily using UCSF ChimeraX version 1.8 (https://www.rbvi.ucsf.edu/chimerax), or PyMOL (https://www.pymol.org/). We typically visually evaluated structure matches for proteins with reasonable scores: mostly but not exclusively, bitscores > 200 to reference structures. Matches where subfamilies had no relatively high scoring matches to a PDB entry were also evaluated, so long as the bitscores were > ∼50. In some cases, structures were dissected and domains evaluated separately (especially where extra domains were present). Protein functional predictions also made use of domain identification tools (e.g., HMMer, https://www.ebi.ac.uk/Tools/hmmer/search/hmmscan), HMM-based analyses and Pfam profiles, (Mistry et al. 2021). In a few cases where there was a sufficient number of reasonably similar Borg proteins available (ideal for multicopy proteins) and structure predictions had low confidence and/or there was little or not detectably structural similarity, protein structure predictions were generated with AlphaFold2 using an input protein multi-sequence alignment without use of PDB templates. Multisequence alignments were generated using MAFFT (Katoh and Standley 2013). Some proteins were also analyzed using ESMfold (https://esmatlas.com/resources?action=fold) and Dali (http://ekhidna2.biocenter.helsinki.fi/dali/). Structural details, including active site residues, were analyzed by reference to the published literature.

Transmembrane regions of proteins were predicted using DeepTMHMM (Hallgren et al. 2022) and signal peptides were predicted using SignalP6.0 (Teufel et al. 2022).

### Phylogenetic analysis of individual protein sequences

#### Dolichyl phosphate mannose synthase tree

DPMS sequences (subfam0199) from Borgs were aligned using MAFFT (Katoh and Standley 2013). and manually refined.The final alignment contained 717 positions. Phylogenetic analysis was performed using IQ-TREE (Nguyen et al. 2015) with automatic model selection and 1,000 bootstrap replicates. Trees were visualized using Geneious Prime 2024.0.4 (https://www.geneious.com) and annotated in Illustrator.

#### B-family DNA polymerase tree

We manually curated B-family DNA polymerases from Borgs, mini-Borgs, bacteria, archaea, and eukaryotic viruses, combining them with reference sequences from Kazlauskas et al. (2020). Sequences were aligned using MAFFT v7.490 (Katoh and Standley 2013) with the L-INS-i algorithm. The alignment was trimmed using trimAL v1.4.rev15 (Capella-Gutiérrez et al. 2009) with the ‘gappyout’ option. The final alignment of 646 positions was subjected to maximum-likelihood analysis using IQ-TREE v2.1.3 (Nguyen et al. 2015). The LG+C60+R+F model was selected based on the Bayesian Information Criterion (BIC). Branch support was assessed using 1,000 ultrafast bootstrap replicates.

#### deSAMPylse-like metalloprotease tree

The collection of putative deSAMPylse-like metalloproteases was established by combining the sequences from Borgs subfamilies assigned to Group 1 with sequences from the complete cMp *Methanoperedens* genome and related sequence recovered via BLASTP from NCBI. Sequences were aligned using MAFFT-LINSi and trimmed using trimAL with the ‘gappyout’ option, resulting in an alignment length of 171 positions. Phylogenetic analysis was conducted using IQ-TREE with the LG+F+G4 model and 1,000 bootstrap replicates.

### Repeat analysis

A custom script (github.com/rohansachdeva/assembly_repeats) for repeat finding and visualization that was reported previously (Schoelmerich et al. 2023) was modified to report the coding and non-coding localization of repeats. Repeats were considered coding if they are fully contained within a coding region.

### Structure-Based HMM Model Refinement

To attribute functions to previously unannotated proteins, we employed a novel structure-based method for the refinement of profile HMMs. We first used BLASTp to retrieve all protein homologs from Borg genomes. We then aligned these sequences using MAFFT v7.490, and manually curated the alignment by removing unaligned residues to restrict the alignment to the core protein domain. We then used HMMer to create a starting profile HMM which we then used to search the UniprotKB database using HMMsearch (Eddy 2011). Hits from this search were then added to the alignment. We then used AlphaFold 2 to predict the structure of the lowest scoring hit above the inclusion threshold (Tunyasuvunakool et al. 2021), and then searched the structure against AlphafoldDB and the PDB using FoldSeek (van Kempen et al. 2023). We added hits from this search to the alignment which we then used to create a new profile HMM. This process was repeated until no new sequences could be added to the alignment. We then analyzed the functional annotations of sequences in the alignment and assigned putative functions to the original proteins.

### Transcriptomic analysis

Total RNA was extracted from ∼5 g wetland soil samples using the Qiagen RNeasy PowerSoil Total RNA Kit. Ribosomal RNA (rRNA) was depleted using the NEBNext rRNA Depletion Kit, followed by a library construction using NEBNext Ultra II Directional RNA Library Prep Kit. The RNA library was sequenced on the Illumina NovaSeq6000 PE150 platform in Maryland Genomics to generate metatranscriptomic data.

Raw sequencing data was processed using BBDuk (https://jgi.doe.gov/data-and-tools/software-tools/bbtools/) to remove low-quality reads. Putative rRNA reads were filtered by SortMeRNA (v4.3.6) (Kopylova et al. 2012). mRNA reads were mapped to genomes using bbmap (https://jgi.doe.gov/data-and-tools/software-tools/bbtools/) with a minimum ident of 97% to calculate the transcriptional activities of genes.

## Reporting Summary

## Data availability

Prior to publication, the data reported in this study can be accessed via XXX

## Author contributions

JFB designed the study. Protein subfamily and detailed protein structure analyses were performed by JFB, with substantial input from GK and specific input from FC (capsid-like proteins) and RB (glycoconjugates). JFB generated the sequence collections with assistance from SL and RS and input from FA, ZB, and LVA. GK performed most analyses of histone-associated proteins, IsrB, TnpB, and Cas12f. LVA made the final versions of the phylogenetic trees. CR performed the dN/dS analysis and the structure-informed HMM search to uncover possible phage structural proteins. L-DS generated and analyzed the transcriptomic data, identified putative packaging genes, and refined the synteny analysis for one region by adding Mini-Borg data. Repeat analyses were performed by RS, who wrote the script, and JFB who analyzed the results. JFB prepared the figures and wrote the manuscript, with substantial input from GK and specific input related to *Nucleocytoviricota* by FA, ZB and FC. The manuscript was revised based on comments from all co-authors.

## Acknowledgements

Innumerable people provided input, but specific thanks to Drs. Ben Adler, Brady Cress, Rotem Sorek and David Komander. Dr. Mart Krupovic and Owen Tuck are thanked for thoughtful comments on the manuscript. LVA support was provided by a UC Berkeley dissertation year fellowship. Funding was provided by the Gates Foundation and the Chan Zuckerberg Initiative to JFB. Funding to FOA and ZKB was provided by a National Institutes of Health grant (no. 1R35GM147290-01), to FC by the Australian Research Council (DP21010338, DP230101879), and to GK by the Snow Medical Research Foundation (SMRF2021-276). Snow Medical were not involved in the design of the study, data collection, analysis, interpretation of the data, the writing, or the decision to submit the article for publication.

## SUPPLEMENTARY INFORMATION

### DeSAMPylases

The presence of the nucleophile serine in the active site region distinguishes metalloproteases from cysteine proteases that have the catalytic triad of cysteine and nearby histidine and aspartate residues. Interestingly, a *Mycobacterium ulcerans* protein annotated as a CysO-cysteine peptidase (AF-X8F1Y7) shares the active site residues of the putative de-SAMP metalloproteases and aligns, for example, with Orange 478 and cMp 650 putative deSAMPylases. The *M. ulcerans* protein is likely not a cysteine peptidase. We conclude that deSAMP-like metalloproteases analogous to those of Borgs and *Methanoperedens* also exist in these bacteria.

### Structural and potential functional diversity across DMPS clades

#### Borg and related protein DPMS clades

The putative protein sequences from Borgs (subfam0199), the complete *Methanoperedens* genome (cMp) and the most closely related sequences from NCBI were analyzed phylogenetically. NCBI sequences are from other *Methanoperedens*, some methanogens, Thermococcales, Bathyarchaeia, Thermoplasmatales, some DPANN and unclassified archaea, and many sequences are from Chloroflexi. Chloroflexi proteins have been repeatedly observed to place phylogenetically with those of archaea (e.g., (Hug et al. 2013), and vice versa. The pattern supports extensive gene transfer between Chloroflexi and *Methanoperedens*.

#### Structural variation

Representative sequences from Borg and *Methanoperedens* clades were aligned to each other and 5mm0, as well as to other PDB models for proteins with the same DPMS function. All align closely in the cytoplasmic component of reference PDB structures and some align very closely over the entire protein (**Figure S5A)**. Some sequences completely lack the membrane anchor region (yet are still profiled as DPMS; **Figure S5C**).Flanking genes are generally not good candidates for a separate membrane subunit that could form a complex representing the complete protein. Many AlfaFold DB sequences (e.g., *Methanocaldococcus jannaschii* MJ1222, many bacteria and a sequence from a viral metagenome) also lack the membrane anchor. Although some Borg proteins and a few other proteins have an extra cytoplasmic domain (**Figure S5A**), no analogous sequences were found in public data via Foldseek or searches.

#### Key residues

To predict functional relatedness to the biochemically characterized type-I, -II and -III catalytic DPMS family representatives we searched for 40 expected, conserved residues reported in Supplementary Figure 3 of (Gandini et al. 2017). These are involved in metal binding, catalysis and substrate interactions. Residues were considered comparable if they have biochemical characteristics similar to those identified in the experimental studies. Some proteins, such as Cobalt Borg 211 have all residues as expected based on 5mm0 and the acceptor loops of the Borg protein and 5mm0 are clearly aligned (**Figure S4A**), but most differed in some way from the references. Given the huge phylogenetic spread of proteins under consideration, for the final analysis, we prioritized the DxDxQ of the DADLQ region of 5mm0 that is required for metal binding (particularly, the 2nd D and Q (or similar).

#### Potential functional diversity and evolution

Presence/absence of the membrane anchor and the presence of residues of the expected metal binding motif were mapped onto the phylogenetic tree (**Figure 3**, **Supplementary Data item 4**).

Clade 1 is comprised of 9 Borg proteins that include the anchor region and all have the DxDxQ residues.

Clade 2, sibling to Clade 1, is comprised of sequences from diverse archaea, two sequences from *Methanoperedens*, as well as sequences from Thermoplasmatales, Bathyarchaeia, Chloroflexi, and a single Purple Borg sequence. The proteins possess the cytoplasmic domain and the membrane anchor, with the exception of a sub-clade that includes one Chloroflexi, one *Methanoperedens* and two Borg sequences that lack the membrane anchor. The placement within a much larger clade of proteins with the membrane anchor suggests these anchor-free variants derived from an ancestral protein with an anchor. All Clade 2 sequences have the DxDxQ residues, except one sequence from Purple Borg, which has NxDxQ, and is likely able to bind the metal.

Clades 1 and 2, along with one sequence from Iris Borg and seven sequences from various archaea all have the capacity to bind the metal, and represent a major subdivision in the tree.

Clade 3 is comprised of 6 Borg sequences and a single sequence from the cMp genome. All proteins lack the membrane region and lack the expected glutamine residue. Instead, these proteins have S, N or A. Of these, only those with N instead of Q may bind the metal.

Clade 4 is comprised of 10 Borg sequences, all of which lack the membrane region and all lack the DxDxQ residues, with S in place of Q, as for a subset of sequences in Clade 3.

Clade 5 is comprised of 5 Borg proteins that lack the membrane region and with DxDxT instead of DxDxQ. Closely related is Viridian 228, which features a membrane anchor, has T in place of Q.

Clade 6 is comprised of 8 Borg proteins and one protein from the cMp genome. All possess the membrane region and, like Clade 5, have T in place of Q.

Clade 7 is comprised of numerous database sequences from *Methanoperedens* and other ANME archaea (collapsed), 19 Borg sequences that form a sibling sub-clade and two more distantly related cMp sequences.

Clade 7 also contains two partial sequences from Brown Borg (377 - 378) that represent a split protein. The genome is fully supported by reads, thus this is not an assembly error. The gene switches from +2 to +3 frame and 377 has a short C-terminal extension (**Figure S7**). A +1 frame shift follows a “slippery” polyT (and generally AT-rich) region, as is often the case for frameshifted genes. Assuming frameshifting leads to a complete protein, all Clade 7 representatives have the membrane anchor and the expected DxDxQ motif. Basal to Clade 7 Borg are two Methanoperedens cMp proteins with the membrane anchor and the expected DxDxQ residues.

Clades 3-7 plus a few intervening sequences, represent the second major subdivision in the tree. Sub-clades of proteins without the membrane anchor phylogenetically intersperse with those possessing the membrane anchor. The tree topology is suggestive of evolution from a membrane anchored ancestor (as for the first major subdivision), potentially with three separate membrane anchor loss events with a single origination of the DxDxT substitution.The placement of single *Methanoperedens* proteins in Borg-dominated Clades 2 and 3 suggest that the host acquired these variants via lateral gene transfer from Borgs whereas the reverse pattern in Clade 7 is best explained by gain of Borg proteins from *Methanoperedens* or related archaea.

#### Extra cytoplasmic domain proteins and possible multimerization

The Borg, *Methanoregula* and *Methanoperedens* genomes with extra cytoplasmic domain are consistently profiled as DPMS. The next best annotation (Dali) is glucosyl-3-phosphoglycerate synthase (also has only one cytoplasmic domain) in which the metal binding site is formed by widely separated Asp and His (thus not resembling the DxDxQ metal binding site of DPMS).

Some Borg proteins (e.g., Cobalt and Olive), and proteins identified in *Methanoregula* and a *Methanoperedens* genomes with an extra cytoplasmic domain form homotetramers based on alignment of the hydrophobic regions (**Figure S6B**) with modest confidence. Tetramerization is not unexpected for DPMS itself, as PDB 5EKE is known to tetramerize (and experimental methods would have inhibited detection of multimers in several other structure characterization studies, e.g., 5mm0).

All seven proteins with an extra cytoplasmic domain contain metal binding sites in the main cytoplasmic subunit and in two cases, the DxDxQ motif also occurs in the additional cytoplasmic subunit. In the Cobalt Borg protein, the N in the Q site could enable metal binding. In four cases, the first cytoplasmic domain has D/ExDxR/K, but the activation loops are either not present, or additional loops contribute bulky aromatics to the active site that preclude access of the GDP-Mannose donor substrate, suggesting that they may have a function other than mannose donation to a substrate.

### Gene clusters largely implicated in surface modification

In addition to DPMS, Borgs encode many multicopy genes apparently involved in production of extracellular polymers and glycosylation of extracellular proteins and/or lipids and nucleotide sugar transformations. Prominent are, for example, UDP-sugar-epimerases (subfam1264), including UDP-N-acetylglucosamine 4-epimerase, UDP-Glucuronic acid 4-epimerase, GDP-mannose-3’, 5’ -epimerase. These are structurally similar. For example, the six Cobalt Borg subfam1264 proteins all align to each other (despite slightly different PDB-based annotations) and likely play a role in production of glycoproteins, glycolipids, or proteoglycans. Also prevalent are proteins for dTDP-L-rhamnose biosynthesis (e.g., glucose-1-phosphate thymidyltransferase, subfam2340) and glycosyl/mannosyl transferases (subfam515, subfam1659, subfam1055).

Within the region depicted in **Figure 4**, Cobalt 305 appears to have genes for synthesis of glucosamine-6P, a precursor of N-acetylglucosamine. Nine genes encode proteins that likely perform reactions involving N-acetylglucosamine (GlcNAc), N-acetylmannosamine (ManNAc), uridine diphosphate N-acetylglucosamine (UDP-GlcNAc), UDP-3-oxo-GlcNAc and UDP-N-acetylmannosamine (UDP-ManNAc). For example, Cobalt 286 is predicted to be (UDP-GlcNAc) 2-epimerase, which interconverts UDP-GlcNAc to UDP-ManNAc, and 298 UDP-N-acetylglucosamine 4-epimerase converts UDP-GlcNAc to UDP-GalNAc. 287 is likely chitin deacetylase (chitin is a polymer of GlcNAc). A putative UDP-ManNAc dehydrogenase may provide UDP-N-acetyl-alpha-D-mannosaminouronate for biosynthesis of sialic acid-like polymers. Intriguingly, many genes in this region of the Cobalt Borg genome (some sequentially encoded) appear to encode the glycosyltransferase MshA), one of the most prevalent multicopy proteins (subfam1055, **Figure 1A**) and the first step in mycothiol biosynthesis. Encoded in close proximity to MshA and DPMS are deacetylases that may function as the deacetylase MshB (some have what appear to be accessory carbohydrate-binding domains). The ligase (MshC) and acetylation (MshD) genes were not identified.

Within the Cobalt Borg genome region shown in **Figure 4** is a cluster of four genes (308-311) with structures indicative of biosynthesis of O-methyl phosphoramidate (MeOPN), an unusual bacterial capsular polysaccharide (Taylor et al. 2017). Cobalt 309, likely L-glutamine kinase, catalyzes the ATP-dependent phosphorylation of the amide nitrogen of L-glutamine to form L-glutamine phosphate, the first committed step in O-methyl phosphoramidate (MeOPN) biosynthesis. 308 then may convert CTP + L-glutamine phosphate to CDP-L-glutamine, 310 may convert CDP-L-glutamine to L-glutamate and cytidine diphosphoramidate. 311 has several possible annotations, but the best annotated Foldseek hit (e-27 and with good structural alignment) suggests CTP:phosphoglutamine cytidylyltransferase, which is also involved in the biosynthesis of the O-methyl phosphoramidate (MeOPN). CTP:phosphoglutamine cytidylyltransferase may displace the pyrophosphate from a nucleoside triphosphate by phosphoramidate to generate the nucleoside diphosphoramidate (these enzymes correspond to Cj1418, 1417, 1416 of (Taylor et al. 2017).

Cobalt 308 is also a CTP:phosphoglutamine cytidylyltransferase, but with an N-terminal PEP-mutase domain, which is notable in that this enzyme can form C-P bonds that are the hallmark of phosphonates. Cobalt 319 encodes a protein structurally very similar to a fusion phosphonate-specific cytidylyltransferase and 2-aminoethylphosphonate (AEP) transaminase, thus also likely involved in phosphonate biosynthesis.

The genomes appear to encode many proteins involved in phospholipid biosynthesis. UDP-GlcNAc 3-dehydrogenase (GnnA) is involved in lipid A production (β-(1→6)-linked GlNAc disaccharide (e.g., Cobalt 300 modeled to 7bvj). Cobalt 320 aligns in the membrane portion with a CDP-alcohol phosphotransferase that transfers a substituted phosphate group from a CDP-linked donor to an alcohol acceptor, an essential reaction for phospholipid biosynthesis. The Borg protein has the exactly conserved active site, however it lacks the N-terminal cytosolic domain. Genes in the same genomic region encode proteins with close structural similarity to Lipid II flippases, which translocate Lipid II across the bacterial cytoplasmic membrane (in bacteria, for synthesis of peptidoglycan). Predicted multicopy glycerophosphodiesterases (subfam2067, e.g., Cobalt 222-225) also may be involved in glycerophospholipid metabolism.

#### Histones and Vault proteins

Evidence for Borg encapsulation motivated a search for genes that may organize and package the genomic DNA. Borg genomes encode either single or multiple credible histone proteins (e.g., Green 372), histone remodeling helicases (e.g., Cobalt 739, 742), histone protein methyl/acetyl transferases, and histone protein demethyl/deacetyl transferases (e.g., Green 1225). We also identified one helicase-based protein with an ATP binding site that is known to associate with histones, but this protein has an extra nuclease NucT domain that may call into question a role for this protein in histone loading/unloading or packaging.

The Amethyst Borg genome encodes two adjacent (811, 812; **Figure S14A**) and at least one additional histone-like protein (823), all of which have very similar structures. The adjacent encoded Amethyst histones form a dimer that aligns well with the known histone dimer configuration **(Figure S14B)**.

Histones packaging may be facilitated by post-translational modification of the histone proteins. Acetylation factors in Borg genomes suggest that the histones can be acetylated to active transcription. Also identified was NatD (in 4u9w is bound to H4/H2A peptide and CoA). This is among the most selective N-terminal acetyltransferases (NATs); its only known substrates are histones H4 and H2A. This protein acetylates Arg3 of the eukaryotic histone. The structure is similar and it has the ASP in the expected position to bind Arg3 and a pocket that would accommodate CoA (acetylation donor) to catalyze heterochromatin formation. Amethyst Borg has a small MORC3 CW domain-like protein This zinc finger domain binds to methylated histone protein tails and recruits protein.

Borg genomes often encode what appear to be major vault proteins that were not identified in the host *Methanoperedens* genomes. Multimer predictions indicate that these assemble into portions of the expected barrel-like structures. However, computational limitations preclude *in silico* assembly the many subunits that would be required to fully define the complex. Intriguingly, in some genomes (e.g,. Ruby) the putative vault is encoded prior to a histone and a universal single copy protein predicted to be an exonuclease (ubiquitous single copy gene of subfam0908). The subfam0908 protein is profiled as 1zbh 3’-end specific recognition of histone mRNA stem-loop by 3’-exonuclease, but just has the exonuclease RNA degrading domain. In Ochre, Orange, Green, Purple, Rose, Sky, Amber, Amethyst, Emerald, and Viridian Borg genomes, this exonuclease gene occurs in close proximity to genes encoding either the putative vault or a histone, possibly suggesting connections between their functions.

#### ESCRT proteins for membrane remodeling and tubulin-related proteins

It was once thought that ESCRT-III proteins, which are involved in processes such as multivesicular body formation, plasma membrane repair, nuclear envelope reformation, and viral budding (McCullough et al., 2018), were specific to Eukaryotes. However, ESCRT-III homologs occur in TACK archaea (Lindås et al. 2008; Samson et al. 2008), where they have been implicated in membrane scission during viral budding (Liu et al., 2017), extracellular vesicle formation (Ellen et al. 2009), and cell division (Tarrason Risa et al. 2020). ESCRT-III proteins (and their regulators) are also encoded by the genomes of some Asgard archaea, supporting the proposal that the eukaryotic proteins have an archaeal origin (Spang et al. 2015). ESCRT III-like proteins (as well as some other ESCRT-related proteins) occur in some Borgs (**Figure S15**). These proteins may create an invagination in the host cell lipid membrane during Borg encapsulation if they function in an analogous way to ESCRT proteins in *Nucleocytoviricota*, which have a lipid component within their capsids.

A large single copy gene (encoding a protein of 1057 ± 27 amino acids) occurs in all but one of the 17 Borgs (subfam2487) and encodes a protein profiled as similar to tubulin / FtsZ (4b45), but alignment is only partial, and involves the start and ends of the Borg proteins. Intervening between these regions of the Borg proteins is a long coiled-coil (e.g., Orange 265). This region is also present in some AFDB sequences, e.g., *Halorientalis regularis* AF-A0A1G7LXG2-F1. The *Halorientalis regularis* protein aligns reasonably well with Borg proteins (at the sequence and structure level). There are also a few examples from *Methanoperedens*.

#### An enigmatic, highly multicopy protein

A highly prevalent multicopy subfamily in the Borg genomes, subfam1834 with 126 members, and up to 10 copies per genome, encodes a small protein that is always found at the start and end of the genomes (with related proteins distributed throughout). It has a strongly positively charged surface and a topology that resembles that of the best (but low scoring) PDB match (4pt7), the initiator RepA, and expression of this gene was detected for up to five of the eight Black Borg genes in 50 cm, 80 cm, 90cm and 115 cm soil datasets. Some members of subfam1834 are somewhat similar to transcriptional regulators. However, co-folding with double stranded DNA using AlphaFold 3 did not predict tight association. Modeling using the alignment in place of AF2 templates generated a similar moderate confidence structural prediction and did not uncover new functional clues.

### Supplementary Figures

**Figure S1:**
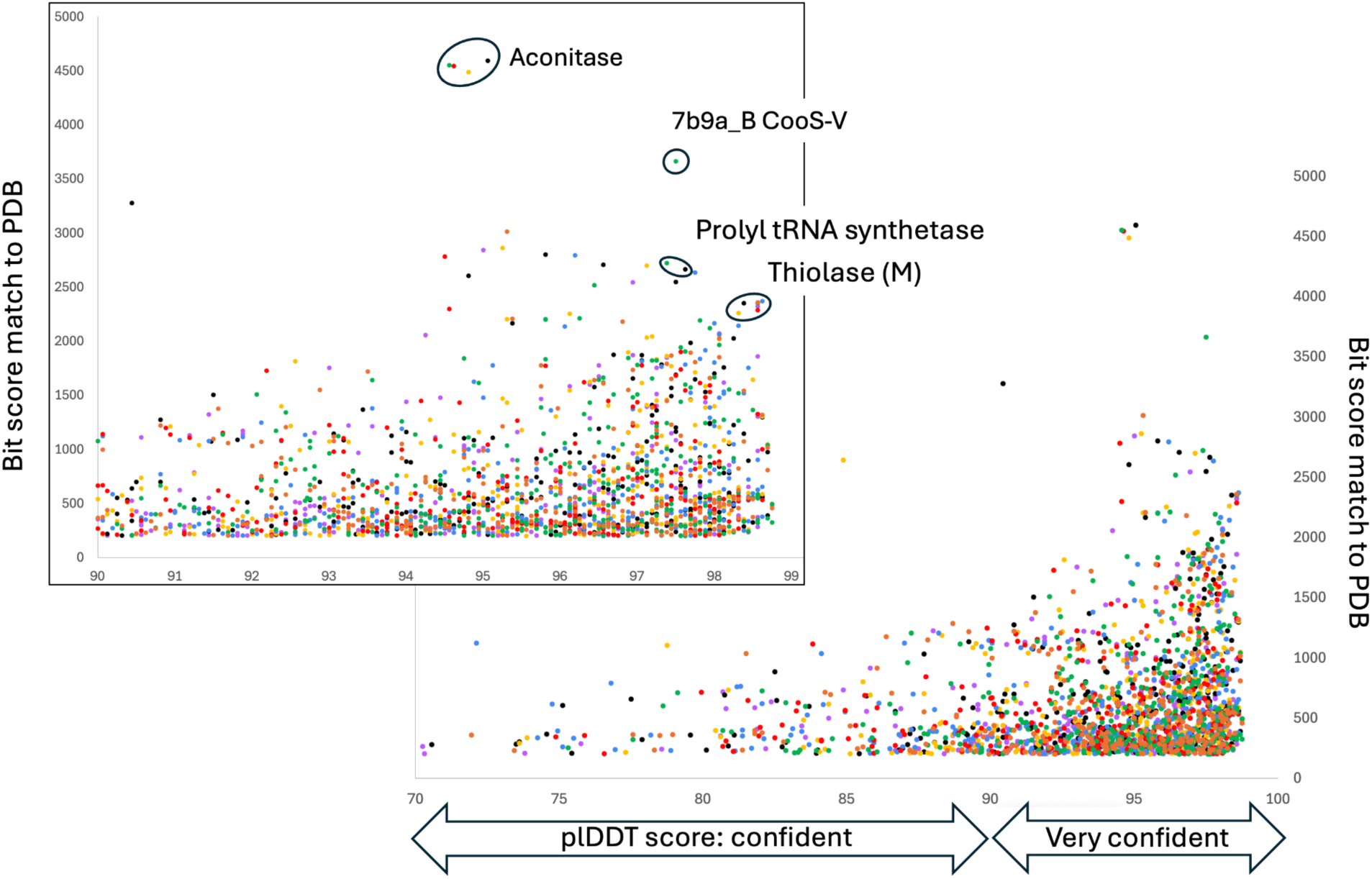
The distribution of scores for proteins with confident (plDDT > 70) or very confident (plDDT > 90) structure predictions and bitscores to best matches in PDB of > 200, colored by Borg of origin. Insert in the upper left shows just the very confident structure predictions and the corresponding bit score of each match in PDB, with some outliers annotated. Calculations for all proteins from 7 complete Borg genomes (9661 proteins) yielded confident or very confident structure predictions in 63% of cases. For tabulation of scores by protein and other statistics, see **Table S1**.

**Figure S2:**
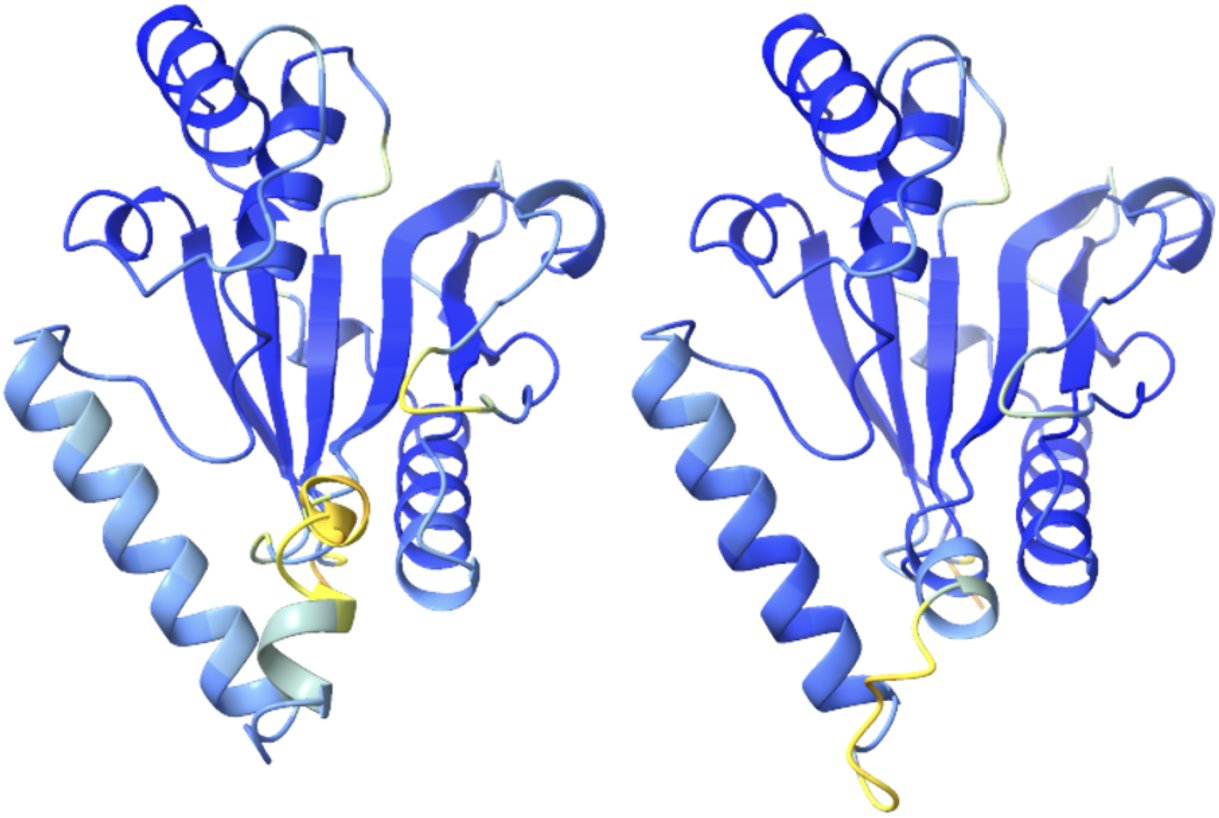
Comparison of folds predicted for Orange Borg 1200, a putative deSAMPylase, with (left) and without (right) use of PDB templates, colored by AlfaFold pLDDT confidence (darker blue indicates higher confidence). The template-free prediction is based on an alignment of the 315 Borg protein sequences (see Figure 1B).The folds have similar fold confidence scores.

**Figure S3:**
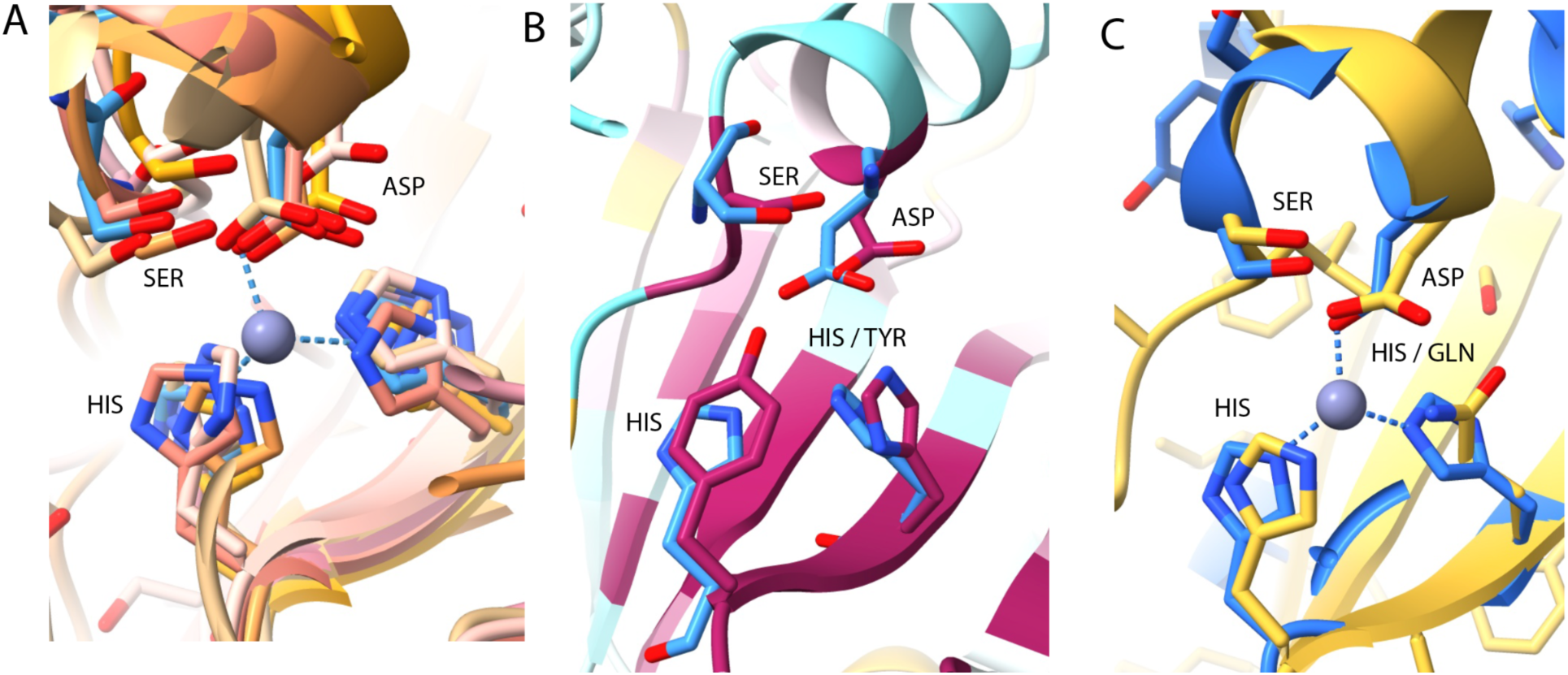
A. Most of the 315 Borg proteins are likely metalloproteases with the HIS,HIS,ASP and SER active site residues, as illustrated by six examples of active site regions of six Cobalt Borg de-SAMP proteins aligned with 3rzu (Crystal Structure of the Catalytic Domain of AMSH), which is shown in blue. **B**. A few Borg proteins do not display these exact expected active site residues, thus may have modified functionality: Amber 1107 colored by conservation (cyan to magenta with magenta being highly conserved based upon 315 sequences), aligned to active site residues of 3rzu (blue). Note the tyrosine in place of histidine in the active site. **C.** Amber 1231 (gold) compared to 4msd (blue), illustrating an active site variant with glutamine in place of histidine.

**Figure S4:**
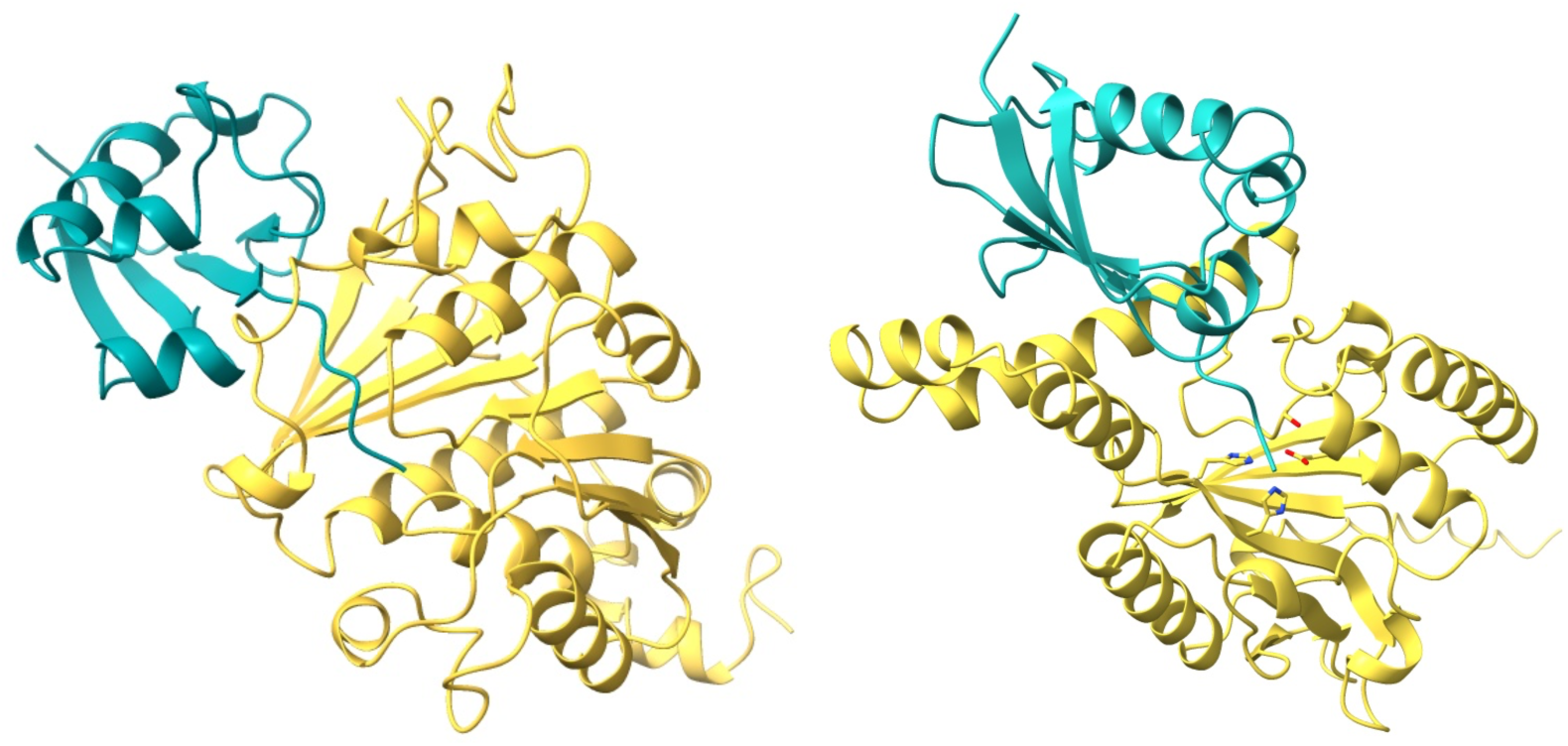
AlphaFold 3 cofolding structure predictions. **A**. The *Methanoperedens* SAMP (cMp 2846, aqua) bound to a *Methanoperedens* SAMP-adding enzyme E1 (cMp 3305, gold), ipTM = 0.86, pTM = 0.92. The multimer involving deSAMP displays the conserved ꞵ-grasp fold and the active site (His,His,Asp,Ser, as shown) engages with the di-glycine tail of the SAMP, as expected. **B.** The structure of the *Methanoperedens* SAMP bound to a putative Borg deSAMPylase (Orange Borg 885), pTM = 0.23pTM = 0.65. The *Methanoperedens* proteins were identified using Haloferax references.

**Figure S5:**
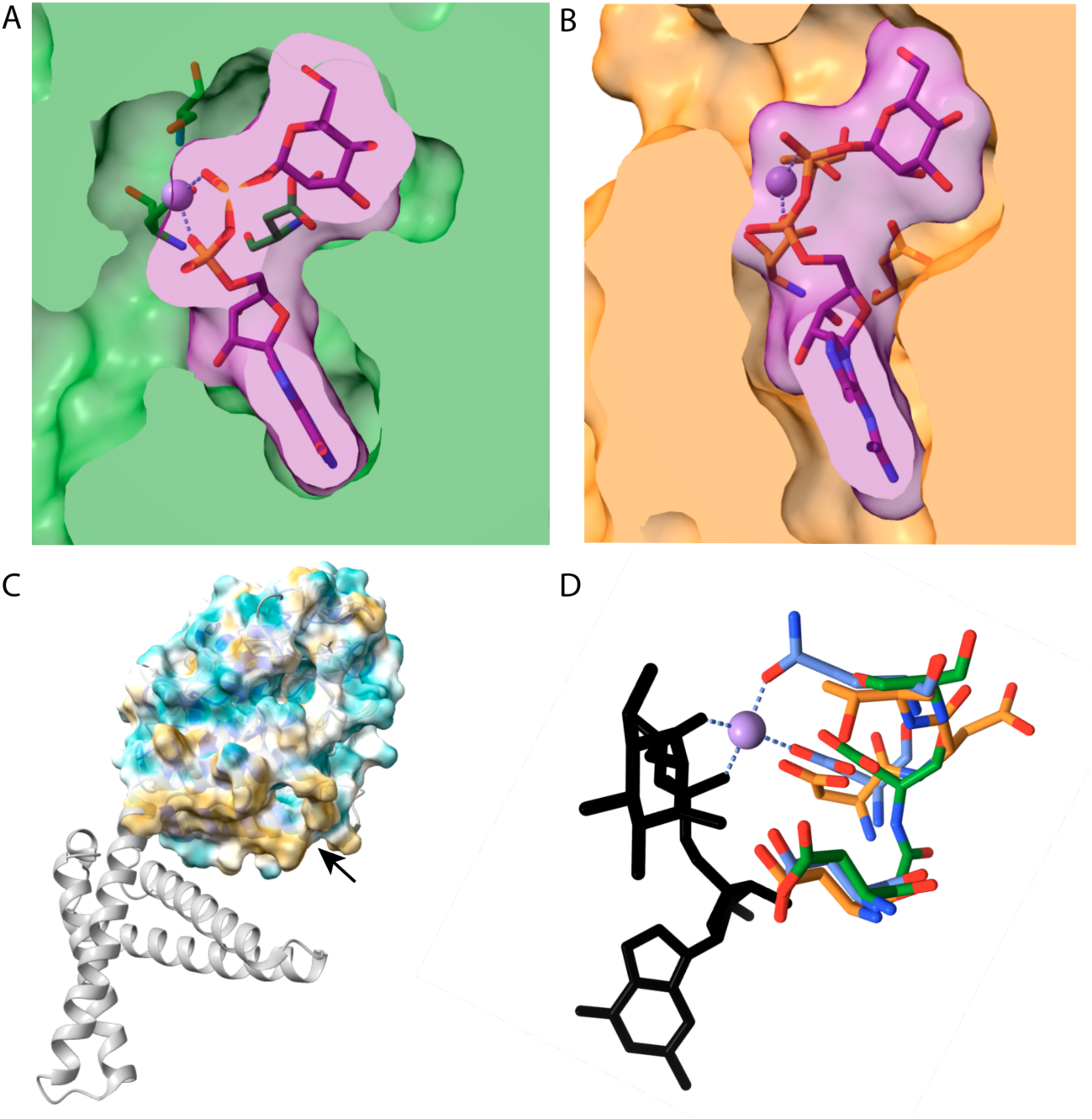
Features of variant DPMS proteins. **A.** Surface representation of the active site pocket of Green 271 reveals a cavity that could accommodate the UDP-Mannose. **B.** The active site pocket of Orange 206 could accommodate the UDP-Mannose. **C.** Anchorless versions of DPMS typically have a hydrophobic patch (gold color, see arrow) in the membrane contact region adjacent to where an anchor occurs in some variants, as shown by 5mm0 (gray). **D.** The conserved D,D and Q active site of 5mm0 (blue) with the Mn shown in purple and the ligand in black, aligned to the residues found in two Borg proteins that lack a membrane anchor and have divergent active sites (Green 271 of Clade 3 (green): S/Q, and Orange 206 of Clade 5 (orange): T/Q, see Figure 3A).

**Figure S6:**
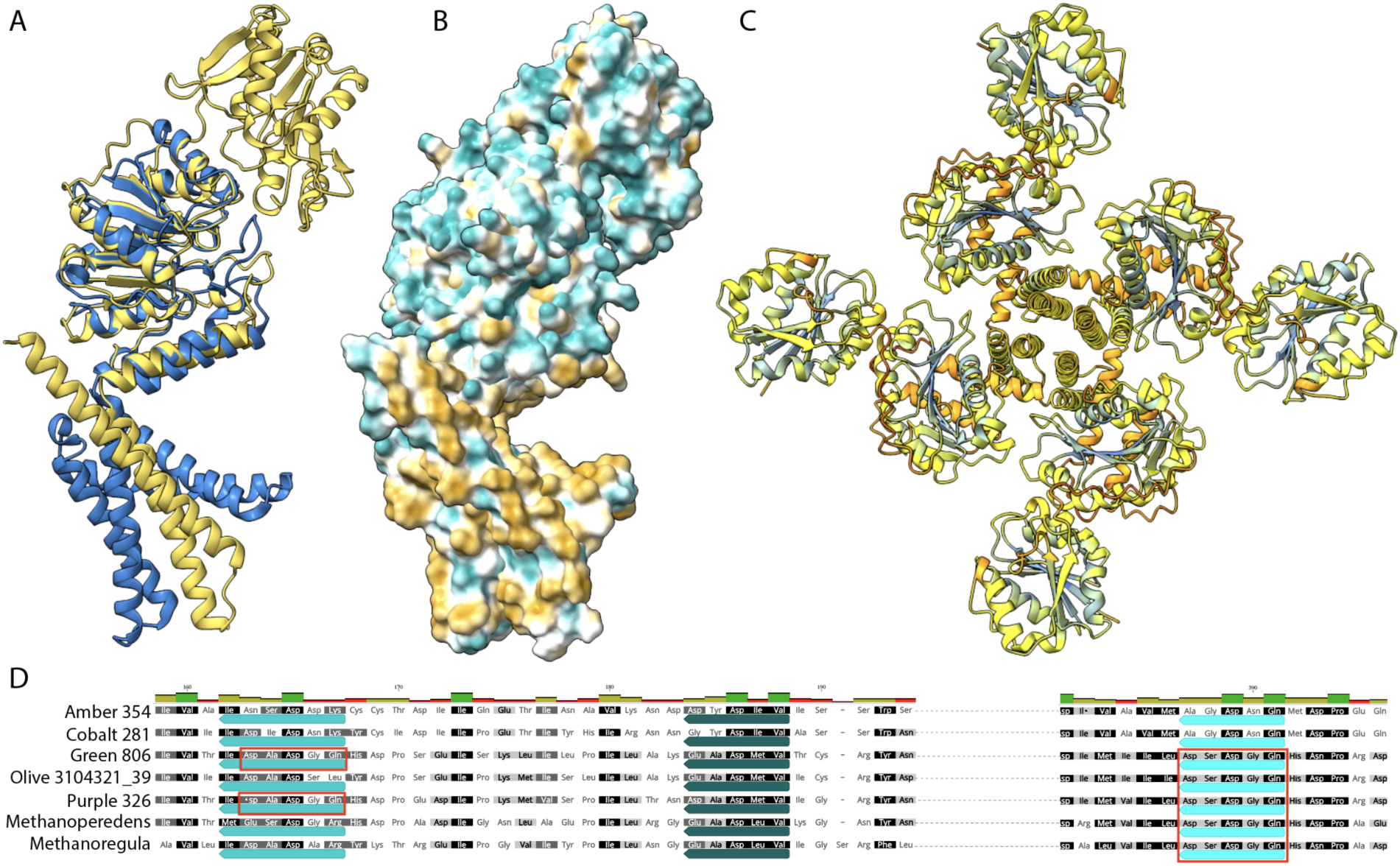
Putative DPMS and variants with an extra cytoplasmic domain. **A**. Green 806 (gold) as an example of proteins with an extra cytoplasmic domain, aligned to 5mm0 (blue) **B.** Hydrophobicity of Green 806, with gold indicating hydrophobic regions that are especially prevalent in the membrane anchor region. **C**. Tetramer of Purple 326 colored by AlphaFold plDDT scores (high confidence is dark blue). Despite modest confidence scores, DPMS tetramer formation is supported by some experimental studies (Ardiccioni et al. 2016). **D.** Regions of the multisequence alignment for examples of DPMS with extra cytoplasmic domain showing the residues of the potentially active site in the main and extra cytoplasmic domain (canonical active sites are indicated by red boxes).

**Figure S7:**
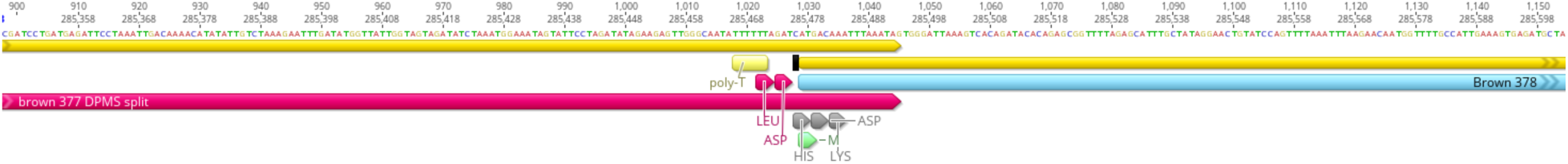
One predicted split DPMS with the DxDxQ motif (Brown 377,378). This should generate a functional protein following a +1 frameshift. The genome is fully supported by reads, thus this is not an assembly error. The gene switches from +2 to +3 frame and 377 has a short C-terminal extension. A +1 frame shift appears to occur after a “slippery” polyT (and generally AT-rich) region, as is often the case for frame shifted genes.

**Figure S8:**
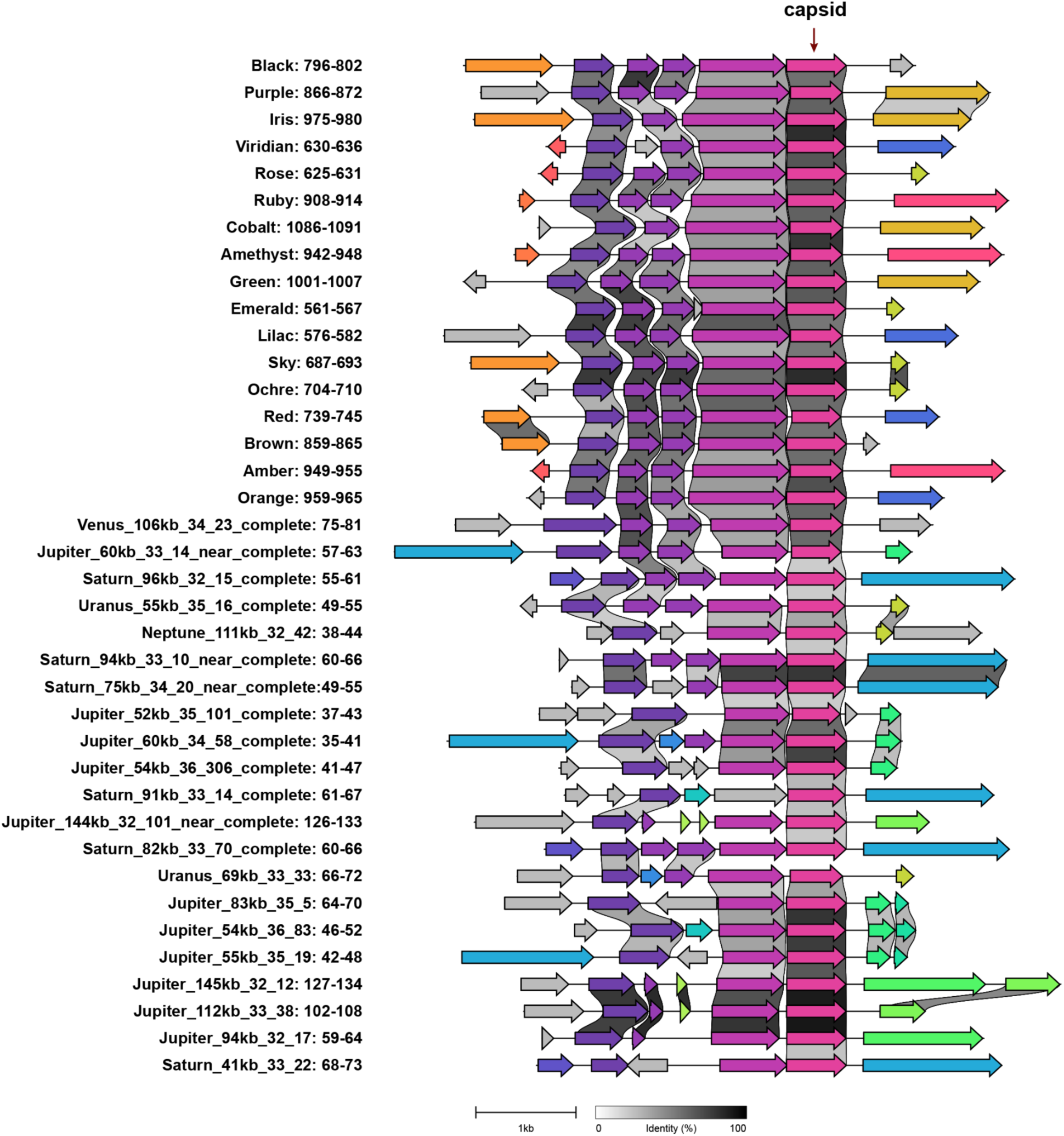
Gene content and gene order conservation in Borg and mini-Borg genomes that encode a gene cluster that features a putative capsid protein (including Black 801), pink. which is highly expressed in some samples, see Figure 5A). The functions of the prior conserved genes could not be discerned by domain analysis, structure prediction analysis (even using the multi protein sequence alignment instead of PDB templates) or multimer calculations. When expressed by Black Borg, the genes in this region are transcribed independently yet when expressed in mini-Borgs, they form a single transcript. Genes classified as homologs share the same color.

**Figure S9:**
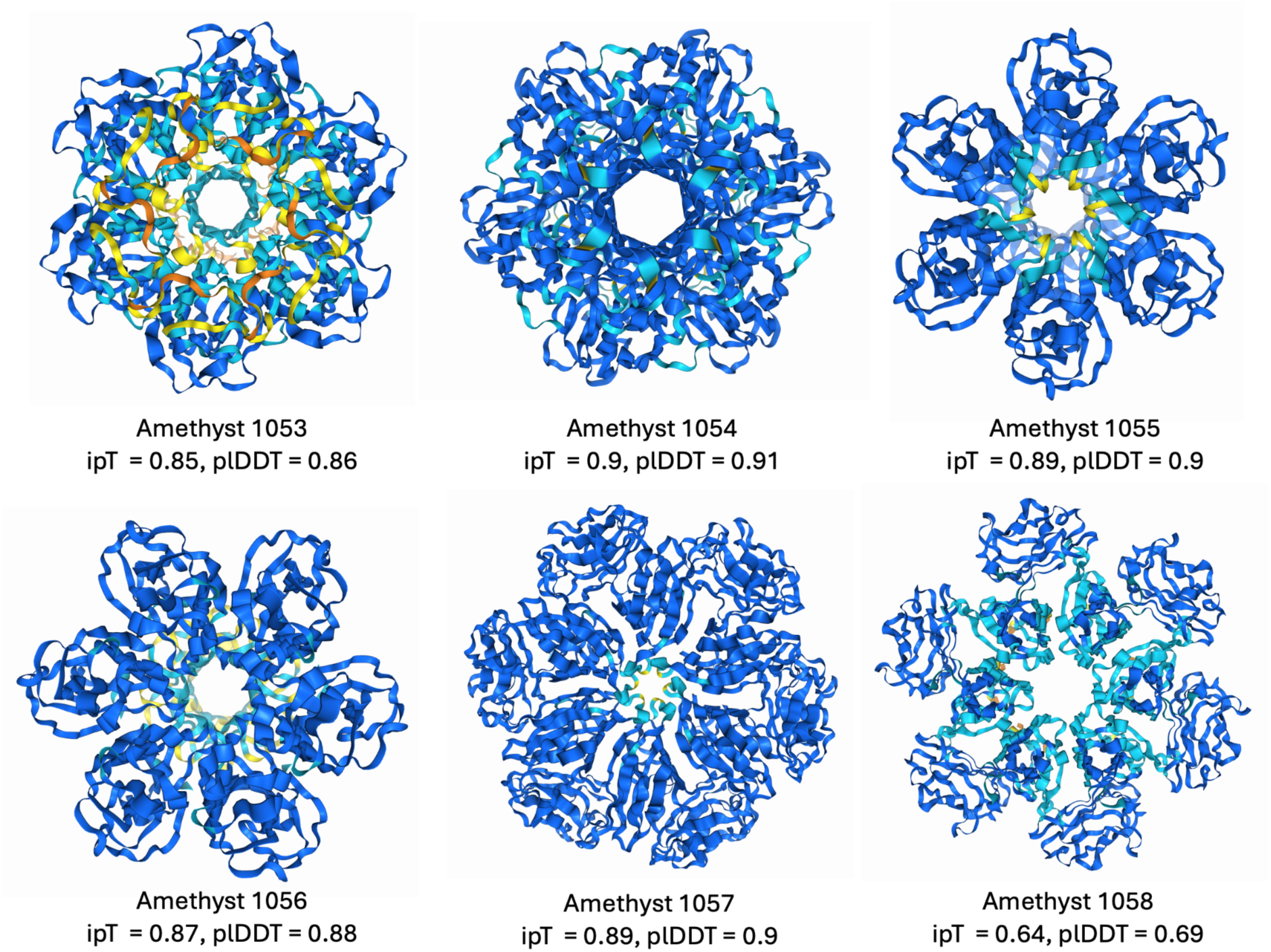
Homo-hexamers of sequential capsid-like proteins from Amethyst Borg (1053-1058). The region encoding these proteins is essentially syntenous in Black, Green, Ochre, Ruby, Orange, Amber, Cobalt, Red and Viridian Borgs. Proteins are coloured by AlphaFold2 pLDDT confidence scores. We also calculated structures for hetero-hexamers of consecutively encoded proteins and found that the multimer confidence scores were lower than for homohexamers (1053 and 1054: pTM = 0.5, pTM = 0.61 and 1055 and 1056: ipTM = 0.58, pTM = 0.63). Thus, we suggest that homohexamer assembly is likely.

**Figure S10.**
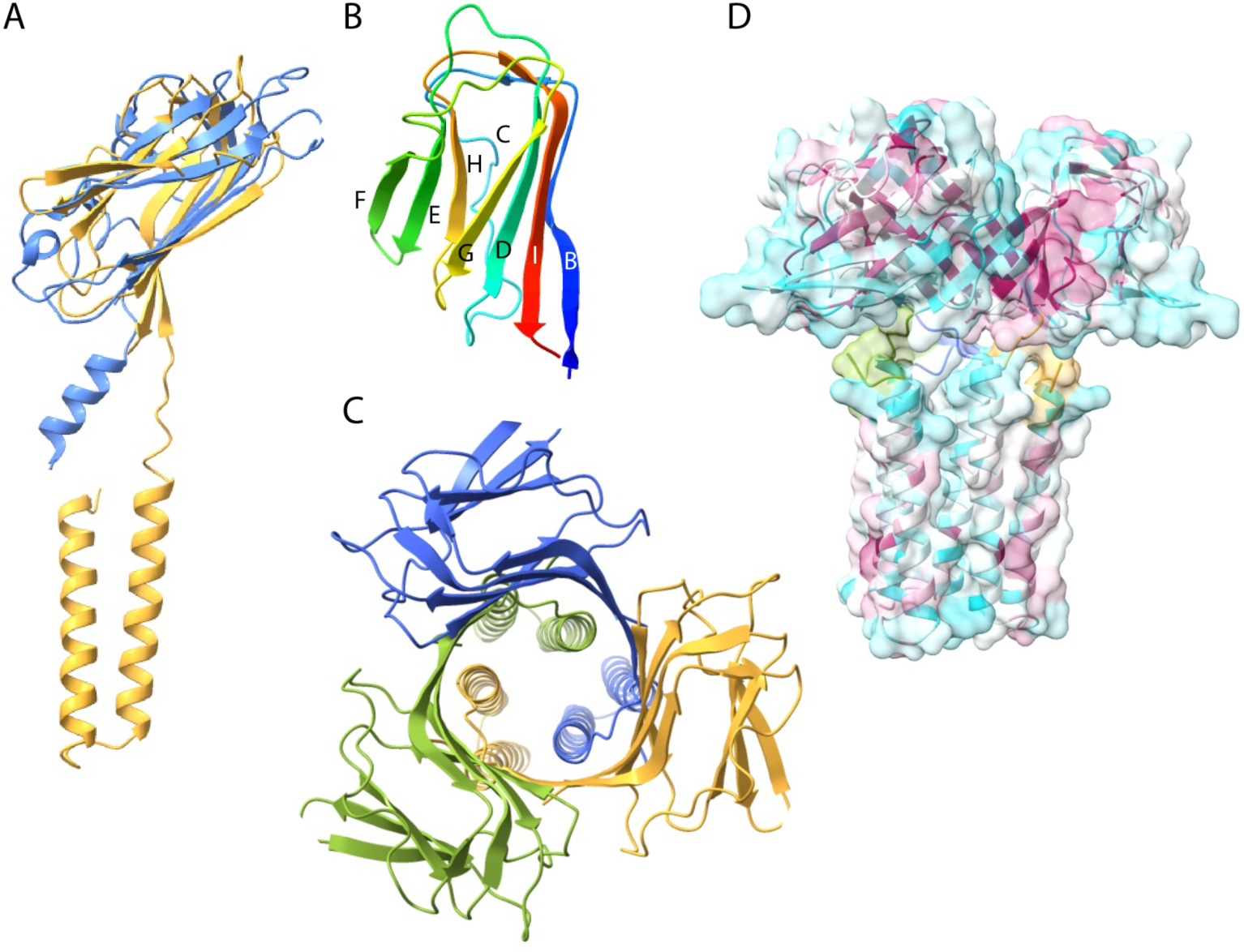
The 17 Borg genomes encode multicopy (subfam1773, 43 total) capsid-like proteins, initially profiled as a bacteriophage tail knob protein (4k6b, a trimer). **A.** Sky 263 monomer (gold) aligned to 4k6b (blue). The alpha helical regions of the Borg proteins are hydrophobic, thus may insert into a membrane. The predicted structure also matches reasonably to the chain B of 6h9c (the capsid-like protein illustrated in Figure 5 aligns with 6h9c VP7 subunit M). **B**. Detail of the beta sheet arrangements in Sky 263 indicating classification as a jelly roll fold. **C**. Sky 263 as a homo-trimer, with each subunit colored differently and viewed from above, first calculation: ipTM 0.67, pTM = 0.71, second calculation ipTM = 0.76. pTM = 0.78. Lowest confidence is localized to the alpha helical regions. Scores for the pentamer are much lower than for the trimer (ipTM = 0.25, pTM = 0.28). **D**. Side view of the Sky 263 homo-trimer colored by conservation (cyan to magenta, with magenta as high confidence) showing conserved residues at the subunit junctions in the jelly roll fold region.

**Figure S11:**
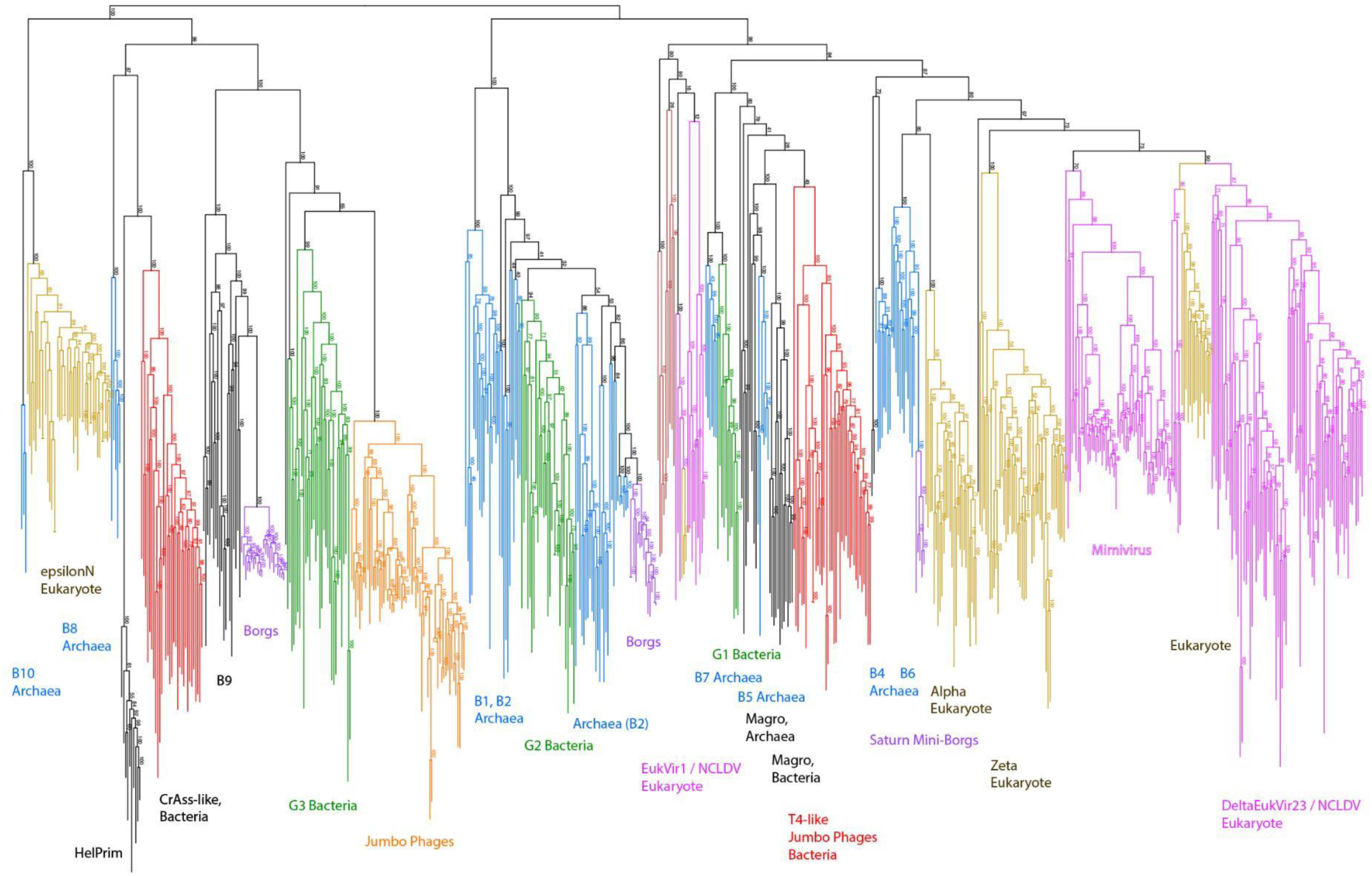
DNA polymerase B tree featuring a sampling of reference sequences (with clade names) from (Kazlauskas et al. 2020) and sequences from internal databases for *Nucleocytoviricota* / mimiviruses, and Jumbo phages. Sequences that are always present in Borgs clade with B9, which Kazlauskas noted are mostly (96%) from metagenomic databases (only two sequences are annotated being from a Thermoplasmatales archaeon and a Candidatus Woesearchaeota archaeon). Other B9 sequences we identified in metagenomic data were likely Woesearchaeota or viral. Other Borg sequences fall into B2, and place with those of host *Methanoperedens* archaea. Mini-Borg sequences fall with archaeal B6. Magro are viruses of marine Euryarchaeota (REF). Borg sequences do not appear to share a recent common ancestor with *Nucleocytoviricota* but rather place firmly with sequences from Archaea.

**Figure S12:**
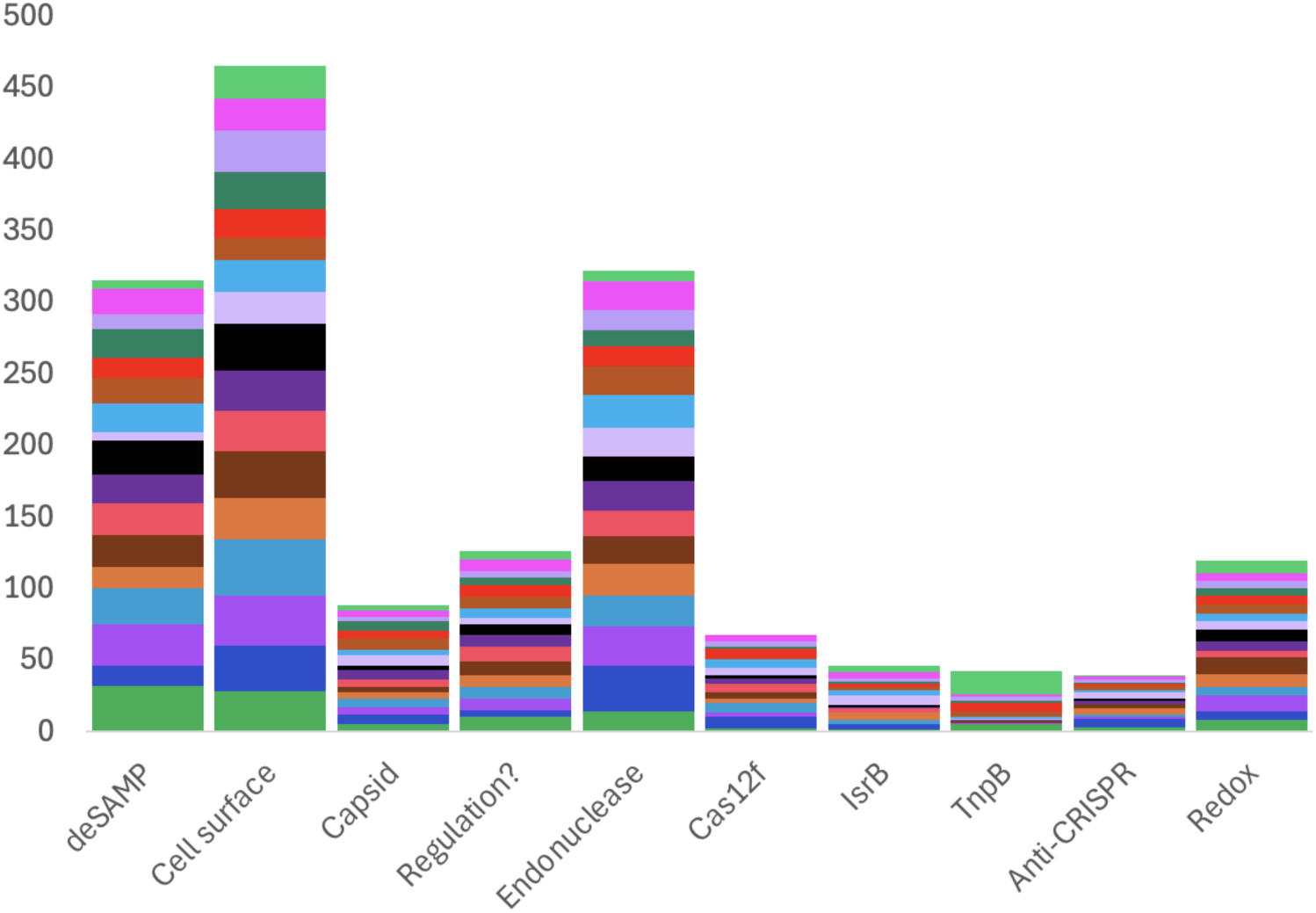
Overview of the more prevalent Borg multicopy proteins (**Table S8**), in some cases groups of subfamilies, into categories based on related functions.

**Figure13:**
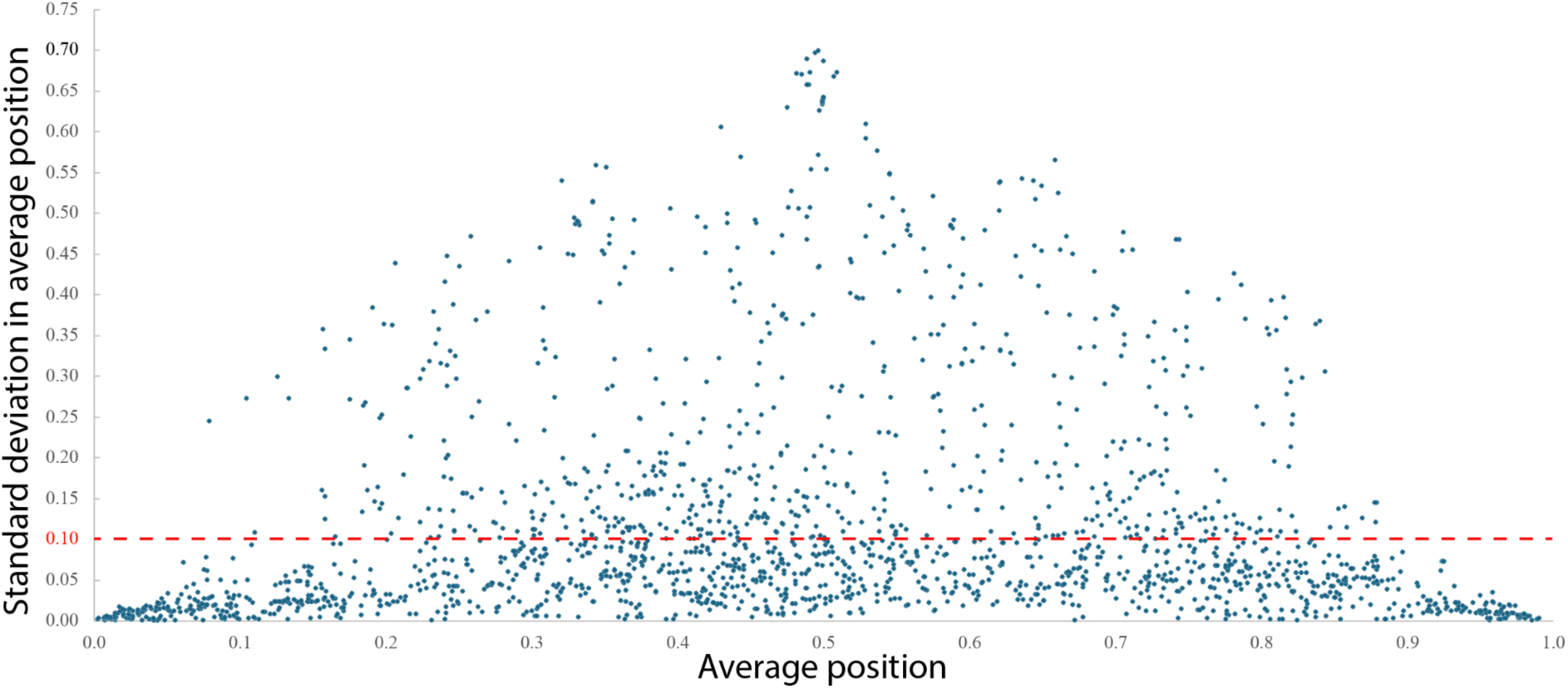
A polt of the average gene position (x-axis) for each subfamily that is not in multicopy in any genome (1931 of 2609 subfamilies) vs. the standard deviation in this position (y-axis). 1243 (64%) of proteins have standard deviation values are ≤ ± 0.1 (dashed red line). 24% of cases with standard deviation values of > ± 0.1 are subfamilies with only two members.

**Figure S14:**
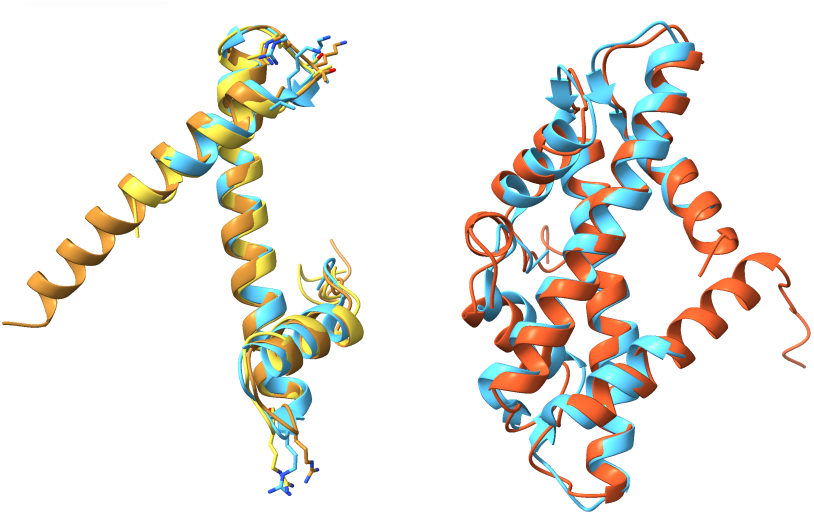
Borg Histones. **A.** Consecutive histone-like proteins from Amethyst Borg are structurally similar (811 and 812, gold and orange), with best match to the experimentally characterized structure of a histone from *Pyrococcus horikoshii* (PDB 1KU5, blue). **B.** Amethyst 811 and 812 form a heterodimer (ipTM = 0.88, pTM = 0.86, red) that aligns well with that of 1KU5 (blue). Related sequences occur in *Methanoperedens* and phylogenetically intermix with Borg sequences.

**Figure S15:**
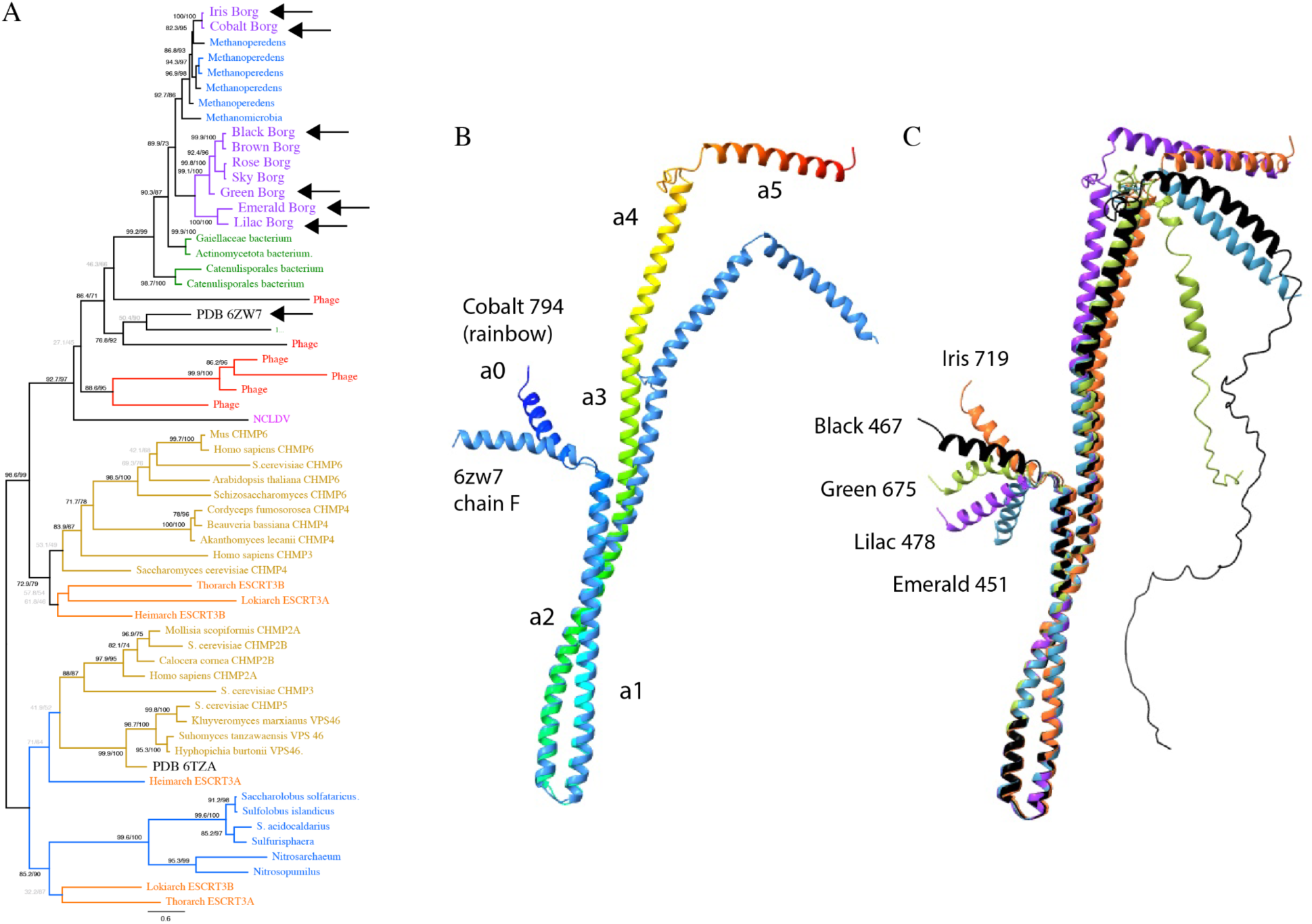
Borg genomes encode membrane remodeling machinery. Among various proteins predicted to have functions analogous to those of the eukaryotic ESCRT system are a subset with structures analogous to that of bacterial membrane remodeling proteins exemplified by PDB 6zw7. These proteins share a common ancestor with the ESCRT III system of Eukaryotes (Spang et al. 2015). **A.** Phylogenetic tree (MAFFT alignment, trimmed using trimal gt 01) including 6zw7, Borg sequences and related sequences from bacteria (green font), archaea other than Asgard (blue font), Asgard archaea (orange font) and Eukaryotes (gold font). Arrows indicate the sequences used in structure predictions in B. and C. Interestingly, the Borg proteins are placed with those from *Methanoperedens* and related archaea, bacteria, and bacteriophages, but not with eukaryotic sequences. **B.** Alignment of the predicted structure of a protein from Cobalt Borg and a subunit of the 6zw7 homo-multimer. The Borg protein is rainbow colored from start (blue) to end (red) and labeled with the region designations analogous to those of ESCRT-III proteins. **C.** A subset of other Borg proteins grouped (based on sequences) into the same subfamily and predicted to have structures analogous to those of 6zw7 and other ESCRT - III proteins.

### Supplementary Tables

**Tables S1 - S7** Listings of the identified the best matches for each protein with a structure predicted using AlphaFold 2 and in the PDB, along with bitscores and e-values, for seven Borgs.

**Table S8** Subfamily counts across the 17 Borg genomes, in some cases grouped by similar functions (blue highlights).

**Table S9** The 17 Borg genomes have one to three genomic regions that encode at least one structural protein (listed by gene number) and occur in a reasonably well-defined region of the genomes. The first region is highly expressed, see **Figure 5**. The proteins in the second region have distant but detectable similarity to bacterial capsid-like / tail-like proteins. The third region features capsid-like proteins featuring a jelly roll fold, encoded consecutively or near-consecutively. These occur in a fairly confined and consistent relative position in each genome. For example, in Green Borg, which has 1517 protein-encoding genes, the three regions occur between gene 1003 and 1101. Genes that were assigned to subfamilies, as noted in different color text: brown = subfam1122, orange = subfam1116, aqua = subfam1011, green = subfam0857, rose pink = subfam2017, blue = subfam1406, purple = subfam1657, red = subfam0634, lime subfam1836.

**Table S10** Detailed version of the **Table 1** overview of the large inventory of genes that are normally only found in organisms and sometimes in giant eukaryotic viruses.

**Table S11** Listing of tandem repeat details for a set of publicly available *Nucleocytoviricota* genomes. Included are columns reporting genome length, GC content, and statistics describing the instances of tandem repeat regions in these genomes. Extracted data (averages, standard deviations) are listed in **Table 2**.

**Table S12** Listing, per *Nucleocytoviricota* genome, of the repeat details statistics including the location of the repeat loci, the lengths of the unit repeats, number of repeat units per locus and the repeat sequence. Extracted data (averages, standard deviations) are listed in **Table 2**.

**Table S13** Listing of tandem repeat details for 17 Borg genomes, including genome length, GC content, and statistics describing the instances of tandem repeat regions in these genomes. Extracted data (averages, standard deviations) are listed in **Table 2**.

**Table S14** Listing, per Borg genome, of the repeat details statistics including the location of the repeat loci, the lengths of the unit repeats, number of repeat units per locus and the repeat sequence. Extracted data (averages, standard deviations) are listed in **Table 2**.

### Supplementary Data items

**Supplementary Data item 1: A.** Predicted structures of all folded proteins for seven Borgs in.pdb format. **B.** Predicted structures for featured proteins from other Borgs, e.g., putative deSAMPs from all Borgs, Sky 263, Apricot packaging ATPas, Purple 326 and Olive Borg, *Methanoperedens, Methanoregula* DPMS proteins with extra domains. **C.** Confidence plots (amino acid specific plDDT scores) for all folded proteins in **A.**

**Supplementary data item 2**: **A.** Phylogenetic tree of putative deSAMPylases showing intermixing proteins from different subfamilies (iqTree, best-fit model: JTTDCMut+F+G4). Nodes with >0.75 bootstrap support are labeled.

**Supplementary data item 3**: Phylogenetic tree of putative DPMS (iqTree, best-fit model: JTTDCMut+F+G4). Nodes with >0.75 bootstrap support are labeled.

**Supplementary data item 4**: Phylogenetic tree for DNA polymerase B (iqTree, best-fit model: JTTDCMut+F+G4). Nodes with >0.75 bootstrap support are labeled.

